# Heterogeneous Exposure and Hotspots for Malaria Vectors at Three Study Sites in Uganda

**DOI:** 10.1101/299529

**Authors:** Su Yun Kang, Katherine E. Battle, Harry S. Gibson, Laura V. Cooper, Kilama Maxwell, Moses Kamya, Steven W. Lindsay, Grant Dorsey, Bryan Greenhouse, Isabel Rodriguez-Barraquer, Robert C. Reiner, David L. Smith, Donal Bisanzio

**Affiliations:** Oxford Big Data Institute, Li Ka Shing Centre for Health Information and Discovery, University of Oxford, Oxford, UK; Department of Veterinary Medicine, Cambridge University, Cambridge, UK; Infectious Diseases Research Collaboration, Kampala, Uganda; Department of Biosciences, Durham University, Durham, UK; Department of Medicine, University of California, San Francisco, USA; Institute for Health Metrics & Evaluation, University of Washington, Seattle, USA; RTI International, Washington DC, USA; Centre for Tropical Diseases, Sacro Cuore-Don Calabria Hospital, Negrar, Verona, Italy

## Abstract

Heterogeneity in malaria transmission has household, temporal, and spatial components. These factors are relevant for improving the efficiency of malaria control by targeting heterogeneity. To quantify variation, we analyzed mosquito counts from entomological surveillance conducted at three study sites in Uganda that varied in malaria transmission intensity. Using a Bayesian zero-inflated negative binomial model, validated via a comprehensive simulation study, we quantified household differences in malaria exposure and examined its spatial distribution. We found that housing quality explained large variation among households in mosquito counts. In each site, there was evidence for hot and cold spots, spatial patterns associated with urbanicity, elevation, or other environmental covariates. We also found some differences in the hotspots in rainy vs. dry seasons or before vs. after control. This work identified methods for quantifying heterogeneity in malaria exposure and offered a critical evaluation of spatially targeting interventions at malaria hotspots.

## Introduction

Despite recent progress in controlling *Plasmodium falciparum* transmission (Bhatt et al., 2015), malaria remains a significant cause of preventable death (WHO, 2016). Human malaria is transmitted by more than 70 species of mosquitoes in the genus *Anopheles* (Service and Townson, 2002). Because of differences in the ecology and the competence of these vectors, malaria transmission intensity is highly heterogeneous over geographies and seasons, and among villages and households (Bejon et al., 2014). Heterogeneous transmission presents both a challenge and an opportunity. Heterogeneous biting propensities among individuals or households (i.e., superspreading) tend to stabilize endemic transmission, but provide and opportunity to increase efficiency of malaria control by targeting interventions at those who are bitten most (Woolhouse et al., 1997; Smith et al., 2007), such as those with the poorest housing quality (Tusting et al., 2017). In countries with low endemic and elimination settings where transmission is maintained in hotspots, spatially targeted interventions are valuable tools in malaria elimination efforts (Bousema et al., 2012).

Geographical patterns, spatial uncertainty, and seasonality in malaria endemicity have been quantified rigorously in recent studies using large aggregated databases describing malaria metrics and environmental covariates (Bhatt et al., 2015; Gething et al., 2016; Cairns et al., 2015), as well as high-quality research data (Guelbéogo et al., 2018; Simmons et al., 2017). Spatial heterogeneity, spatial dynamics, and seasonality are of great interest for spatial and seasonal targeting, which could enable tailoring interventions and coverage targets to the local context and identifying hotspots (Reiner Jr et al., 2015; Ruktanonchai et al., 2016). While these studies capture large-scale spatial and temporal patterns, transmission is a local phenomenon, and many questions about the microepidemiology of malaria remain poorly quantified. Although several studies have also described fine-grained spatial patterns in malaria risk, for example Bejon et al. (2014) and Kigozi et al. (2015), it has proven difficult to quantify heterogeneity in individual or household biting. This is largely due to the fact that the complex endogenous dynamics of vector populations and malaria transmission are often defined by exogenous factors such as local topography, rainfall, hydrology, humidity, temperature, house construction, and malaria control. The prospects for targeting malaria rely on quantifying spatial and temporal heterogeneity and identifying individuals or households with greatest exposure (i.e., hotspots). The scan statistic is commonly used for hotspot identification (Kulldorff, 1997), but the anomalies detected may be stochastic fluctuations that are neither stable nor of any importance for transmission or control. Further, scanning for hotspots does not provide any insight in terms of the drivers of heterogeneity. Accurate quantification of heterogeneity and hotspots at a fine-grain resolution therefore requires taking a different approach.

Understanding malaria transmission and quantifying the possible efficiency gains from targeting interventions require methods to accurately measure heterogeneous biting and its underlying causes. A large longitudinal study of malaria recently conducted in three study sites in Uganda provides a unique opportunity to do so. A notable feature of this dataset is that entomological data were collected longitudinally on all households, so that it was possible to measure and study the household component of biting over seasons and over time. This involves using mosquito count data from monthly entomological surveillance conducted at 330 households between October 2011 and March 2015 for Walukuba subcounty, Jinja District and Kihihi subcounty, Kanungu District; and between October 2011 and September 2016 for Nagongera subcounty, Tororo District.

With the aid of a comprehensive simulation study, we extend the existing methodology in the literature for understanding among-household heterogeneity in malaria exposure and potentially for other mosquito-borne diseases. We evaluated heterogeneous biting propensities among households during different seasons, before and after the application of vector control interventions, and among these three sites which differed substantially in their average malaria transmission intensity. We estimated household biting propensities, which measure the average ratio of mosquitoes caught in a household compared with the population expectation, which is attributable to household characteristics such as housing structure and human hosts within the household. Studies have shown that households within a settlement exhibit spatial heterogeneity in vector distribution, where mosquito densities vary from household to household (Lindsay et al., 1995). It has also been shown that households located close to aquatic habitats tend to receive more mosquito bites due to the relatively higher mosquito abundance in the surroundings and this is known to be affected by wind direction (Midega et al., 2012) and the type of materials used to build the house (Wanzirah et al., 2015). Also, differences in biting attractiveness of human hosts contribute to the variability in mosquito biting among people living in the same household (Lindsay et al., 1993). Individuals who receive the most bites are most likely to be infected and can, therefore, intensify transmission by transmitting malaria parasites to a large number of mosquitoes (Guelbéogo et al., 2018).

Here, we describe the household biting propensities or attractiveness by first estimating and removing the seasonality effects from the household mosquito counts, leaving only the remaining household-level heterogeneity for further inference and study. Next, using these biting propensities, we introduce a novel approach for identifying changes in malaria hotspots over time in which we compute the Getis-Ord statistic (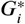) on ratios of household biting propensities for different scenarios, which reflects the relative attractiveness of households in attracting mosquitoes under different circumstances.

## Results

### Quantifying heterogeneous household biting in malaria exposure

A simulation study was designed to determine the most robust and best-fitted model for quantifying household-level heterogeneity in malaria exposure. Due to the presence of excess zeros in the Ugandan mosquito count data (a common feature of malaria vector data), a range of models capable of handling excess zeros were considered, including zero-inflated Poisson, zero-inflated negative binomial, Poisson hurdle, and negative binomial hurdle regression models. The details and the discussion of the results of the simulation study are described in Supplementary File 2. Using pseudo-datasets that mimicked the Ugandan data and several model selection criteria, we found the zero-inflated negative binomial (ZINB) model to be the most robust and best-fitted method for estimating household biting propensities (*ω*), seasonality (*S*), and environmental and measurement noise (*e*) for each study site (Figure 1).

**Figure 1.**
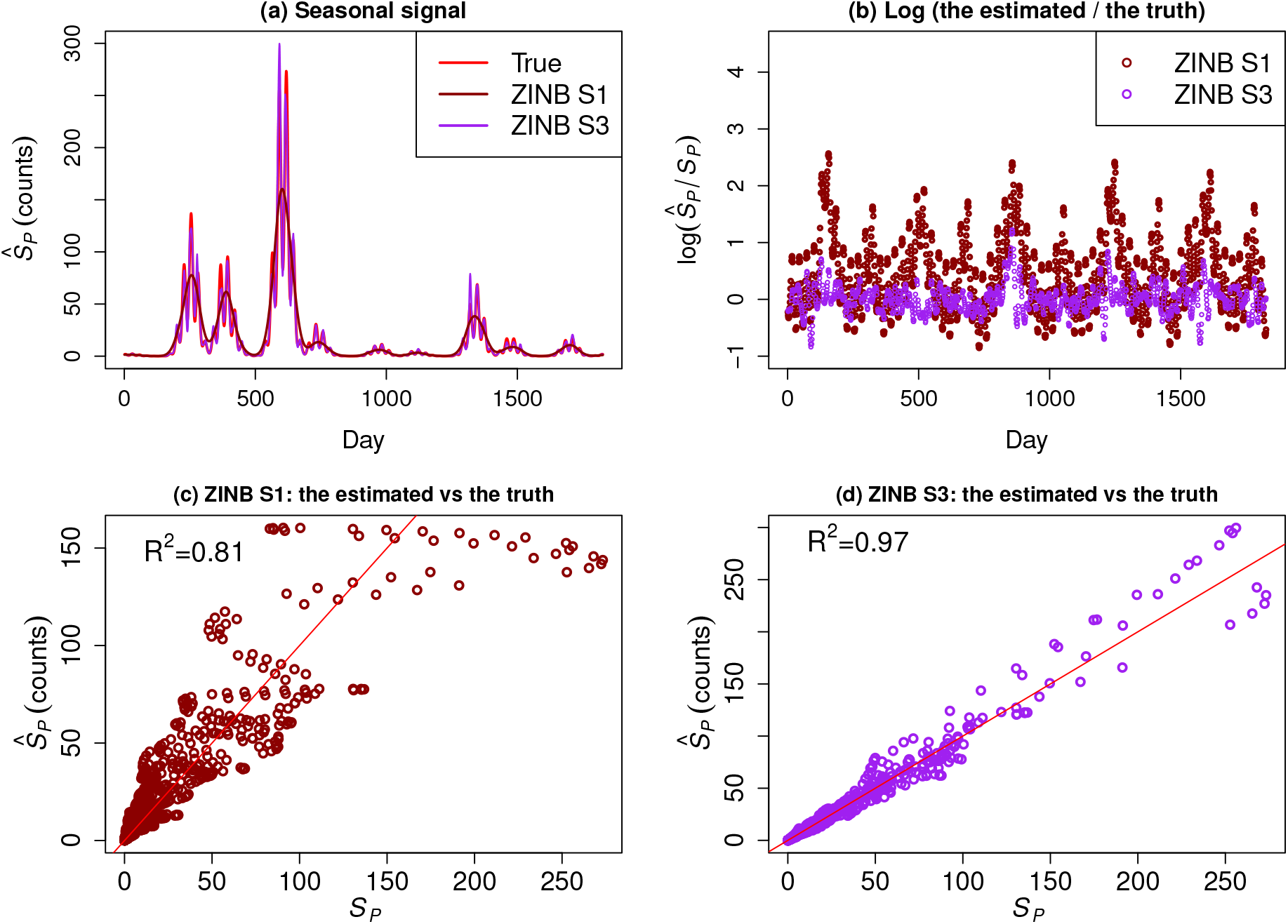
Simulation study: (a) The estimates of seasonal signal (*Ŝ*_p_) for pseudo-dataset D3 reconstructed using the best-fitted model, i.e. the zero-inflated negative binomial model. Smoothing technique S1 (a Gaussian kernel was used to smooth the counts prior to model fitting) produced the worst fit while smoothing technique S3 (a second-order random walk prior distribution imposed on temporal random effects) produced the best fit. (b) The scatter plot shows log(*Ŝ*_p_/*Ŝ*_p_): ZINB S3 resulted in a much better fit of *Ŝ*_p_ than ZINB S1. (c) The ZINB S1 model produced the worst fit when *Ŝ*_p_ (estimated) is fitted against *Ŝ*_p_ (the truth) on a simple linear regression; coefficient of determination, *R*^2^, is around 80%. (d) The ZINB S3 model produced the best fit when *Ŝ*_p_ is fitted against *Ŝ*_p_ on a simple linear regression; *R*^2^ is close to 1.

The overall household biting propensities for the entire duration of surveillance were estimated for each of the study sites, denoted *ω*_overall_. We found patterns among households as well as spatial patterns at a larger scale. Although household biting propensities are closely correlated (by design) with the average number of mosquitoes caught in a light trap per household per night at each study site, they are not identical (Figure 2). Households receiving a larger amount of mosquito bites tend to have a larger biting propensity but this is not always true due to other factors such as seasonality and the environment. A large variation in biting propensities was observed in households across the three study sites, as shown in panel (a) of Figures 3, 4, and 5 for Jinja, Kanungu, and Tororo, respectively. Households with the largest and smallest *ω*_overall_ in Jinja appeared to be located away from the centre of the study region (Figure 3(a)), which is highly urbanized (Kigozi et al., 2015). Households at low elevations in Kanungu (northern part of the study region) generally had a larger *ω*_overall_ while households at high elevations and less rural part of Kanungu (southeast part of the study region) had a much smaller *ω*_overall_ (Figure 4(a)). In Tororo, with the highest transmission intensity among the three sites, some of the households with the largest and smallest *ω*_overall_ seemed to be located around the border of the study region (Figure 5(a)).

**Figure 2.**
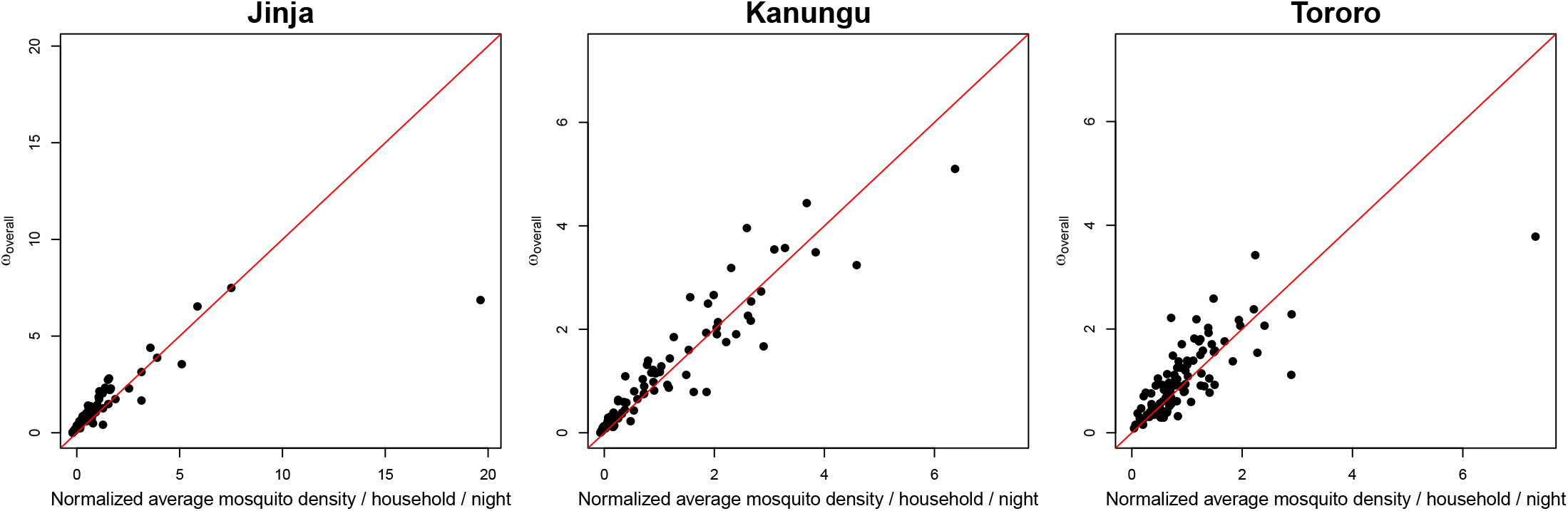
Household biting propensities for the entire duration of surveillance (*ω*_overall_) are plotted against the normalized average mosquito density per household per night for each of the study sites. The red line denotes the 1:1 line. Evidently, *ω*_overall_ had a linear relationship with mosquito density at each study site.

**Figure 3.**
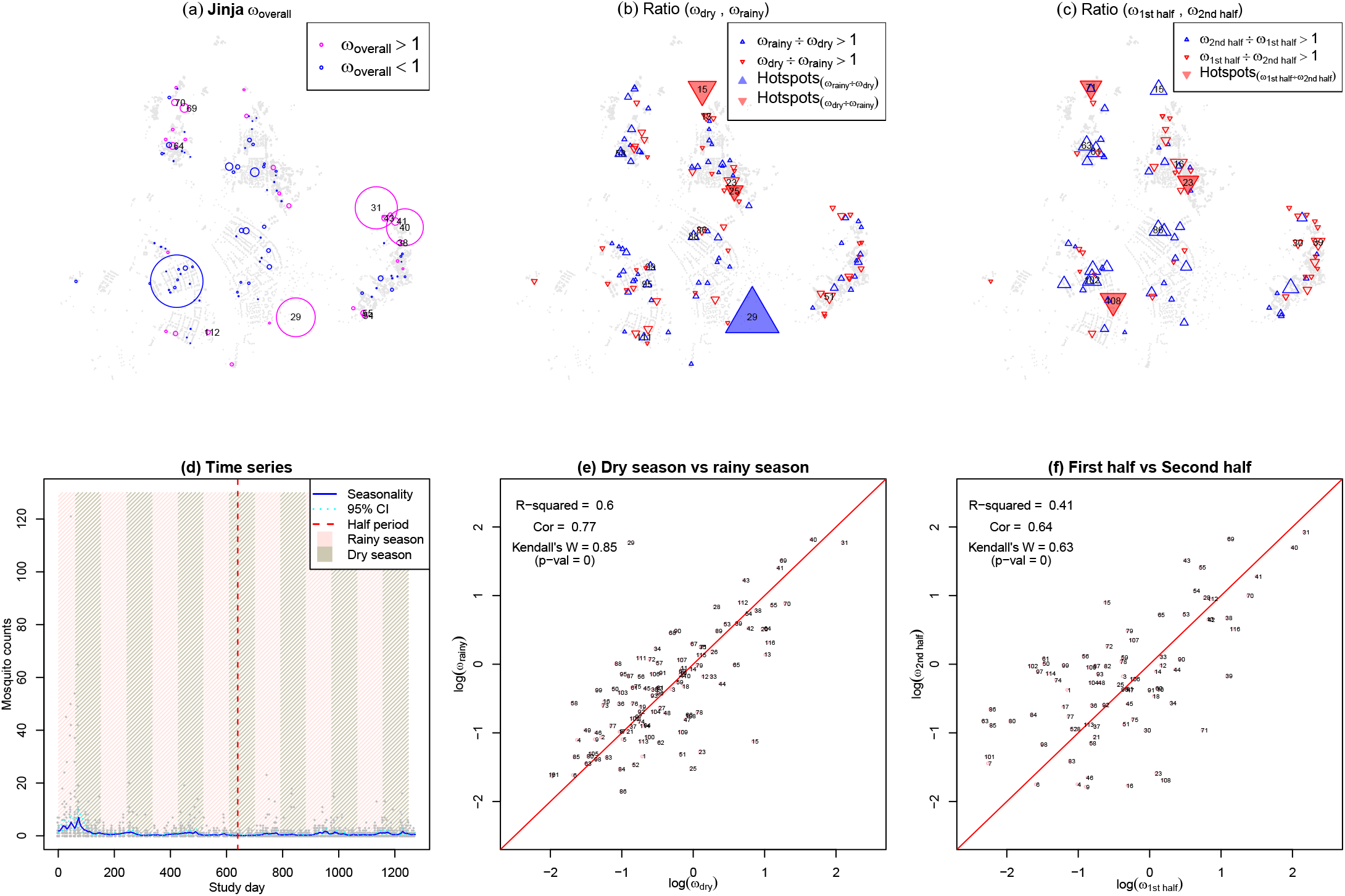
Jinja. **(a)**: Household biting propensities for the entire duration of surveillance. Pink circles denote households with *ω*_overall_ > 1 with the size of the circles showing the magnitude of *ω*_overall_; blue circles denote households with *ω*_overall_ < 1 with the size of the circles showing the magnitude of 1/(3 × *ω*_overall_) to make their sizes comparable to the pink circles; 10% of households with the largest *ω*_overall_ are labelled. The largest pink circle indicates a household with the largest *ω*_overall_ while the largest blue circle indicates a household with the smallest *ω*_overall_. Households with the largest *ω*_overall_ include HH31, HH29, and HH40. Grey dots in the background denote all enumerated households atWalukuba subcounty, Jinja District. **(b)** Ratios of household biting propensities during dry (*ω*_dry_) and rainy seasons (*ω*_rainy_). Each plotted upward pointing blue triangle represents a household with a larger *ω* during rainy season compared to dry season; a blue filled triangle denotes a hotspot for the ratio of *ω*_rainy_/*ω*_dry_. Each plotted downward pointing red triangle represents a household with a larger *ω* during dry season compared to rainy season; a red filled triangle denotes a hotspot for the ratio of *ω*_dry_/*ω*_rainy_. **(c)** Ratios of household biting propensities in the first half period of surveillance (*ω*ı_st half_) and the second half period (*ω*_2ndhalf_). Each plotted upward pointing blue triangle represents a household with a larger *ω* in the second half period of surveillance compared to the first half period; there were no hotspots for the ratio of *ω*_2ndhalf_/*ω*_1st half_. Each plotted downward pointing red triangle represents a household with a larger *ω* during the first half period of surveillance compared to the second half period; a red filled triangle denotes a hotspot for the ratio of *ω*_1st half_/*ω*_2ndhalf_. **(d)** Time series of mosquito count data for the entire duration of surveillance, where each grey circle denotes an observation of a household. The blue solid line denotes the estimated seasonal signal and cyan dashed lines denote the 95% Bayesian credible interval for the seasonal signal. The red vertical dashed line denotes the cut-off for the first half and the second half periods. Rainy season is highlighted in pinkand dry season is highlighted in light green. A weak seasonal signal was observed in Jinja. **(e)** Scatter plot of log(*ω*_dry_) and log(*ω*_rainy_) along with measures of *R*^2^, correlation, and Kendall’s W. For ease of interpretation, we plotted the *ω* on the logarithmic scale. The log transformation also preserved the order of the observations while making outliers less extreme. **(f)** Scatter plot of log(*ω*_1st half_) and log(*ω*_2nd half_) along with measures of *R*^2^, correlation, and Kendall’s W.

### Seasonally varying hotspots of malaria risk

We quantified differences in household biting propensities due to different seasons and vector control interventions using Kendall’s coefficient of concordance (or Kendall’s *W*) (Field, 2005). There are typically two rainy seasons in Uganda (March to May and August to October) with annual rainfall of 1,000 - 1,500 mm. Residents of all study households were provided long lasting insecticidal bednets (LLINs) at enrollment. Over the course of the study compliance (defined as sleeping under an LLIN the previous night at the time of each clinic visit) was over 98% for all three sites. Thus LLIN coverage within the study households (where mosquitoes were collected) was very high and consistent over time and across the three sites. In terms of LLIN coverage in the surrounding communities, coverage levels varied across the sites and over time based on repeated crosssectional surveys. Changes in coverage over time were primarily due to a national LLIN distribution campaign conducted in November 2013 at two sites (Jinja and Tororo) and June 2014 at the third site (Kanungu). However, in a separate time series analysis (Katureebe et al., 2016), there was no significant difference in human biting rates (mosquito abundance) before and after community LLIN distribution for all three sites. Indoor residual spraying of insecticide (IRS) was only used in Tororo, where 3 rounds of the carbamate insecticide Bendiocarb were initiated in December 2014, June 2015, and December 2015 in Nagongera.

To study the impact of different seasons on household biting, we explored biting propensities during dry (*ω*_4y_) and rainy seasons (*ω*_rainy_) for all three sites. For Jinja and Kanungu, we evaluated changes in transmission in the first and last half of the study by estimating biting propensities in the first half period of surveillance (*ω*_1st half_) and the second half period (*ω*_2nd half_). For Tororo, the impact of IRS on household mosquito biting was examined by computing biting propensities before the deployment of IRS (*ω*_before_ įr_S_) and after (*ω*_after IRS_). In Figure S1, we plotted *ω* for different scenarios against *ω*_overall_, changes of transmission over time was confirmed in Jinja while the application of IRS in Tororo had affected the *ω*.

Using the 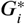 statistic, we found seasonal varying hotspots of malaria risk at the household level for different scenarios at the three sites. Here, a seasonal varying malaria hotspot refers to a household (HH) with drastic differences in *ω* under different scenarios and was surrounded by other households with less drastic changes in *ω*. In Jinja, three households showed considerable differences during dry and rainy seasons while three households were identified as seasonal varying hotspots when compared in the first and last half of the study (Figure 3(b-c)). Being the study site with the lowest malaria transmission intensity, Jinja exhibited a weak seasonal signal in mosquito counts over time (Figure 3(d)). It is noteworthy that HH29, located next to a swampy area near Lake Victoria, had a much larger *ω* during rainy season compared to dry season. Most households in Jinja behaved similarly during dry and rainy seasons, whereas greater differences in *ω* were shown between the two time periods (Figure 3(e-f)).

In Kanungu, there was a seasonal varying hotspot for the dry vs. rainy seasons comparison while three households were identified as seasonal varying hotspots when compared in the first and second half of the study (Figure 4(b-c)). Kanungu was the study site with intermediate transmission intensity and thus exhibited a moderate seasonal signal in mosquito counts over time (Figure 4(d)). Most households in Kanungu behaved very similarly during different seasons and time periods despite some slight differences in the first half and the second half periods of surveillance (Figure 4(e-f)).

**Figure 4.**
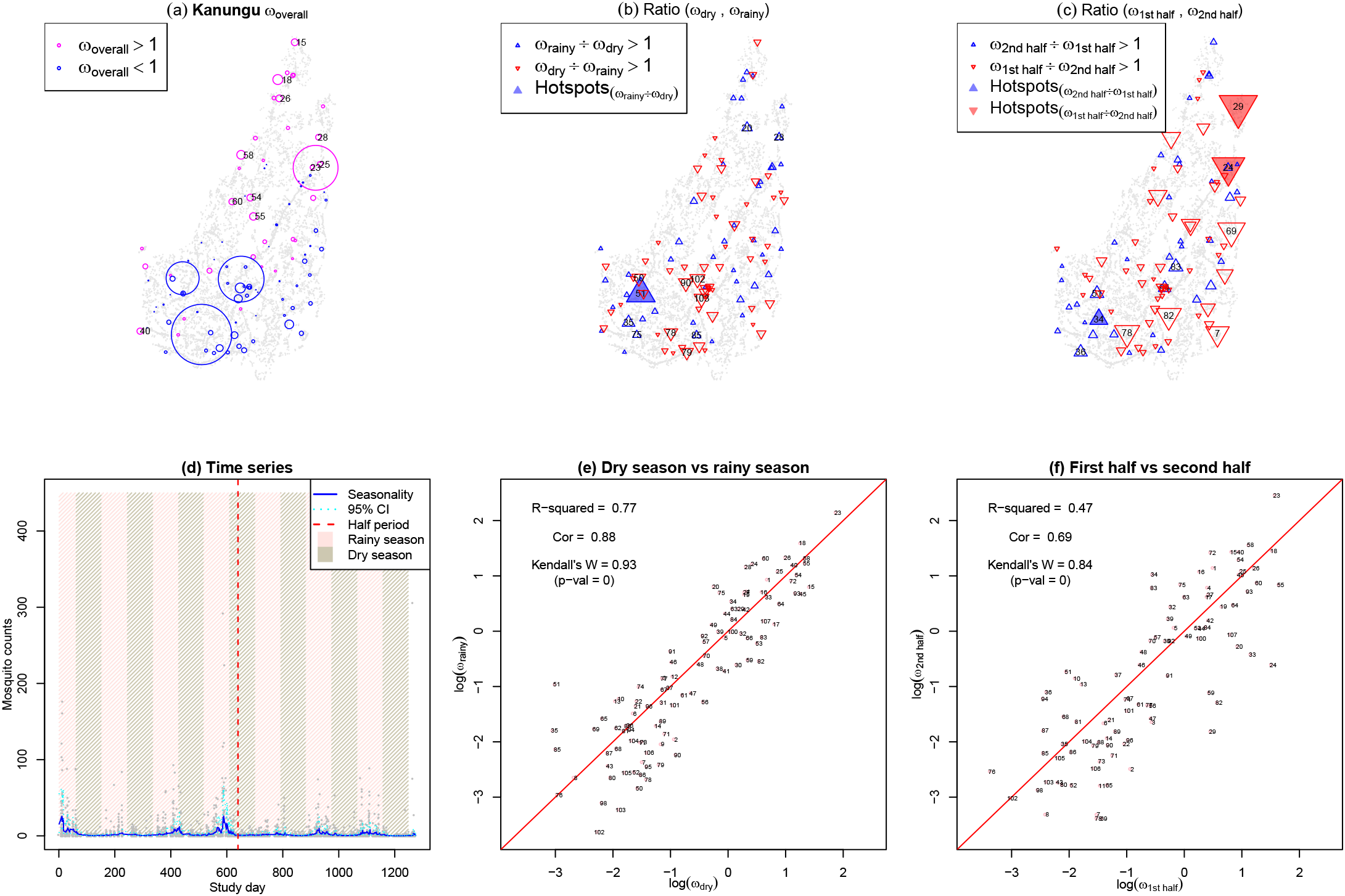
Kanungu. The layout of the figure is the same as in Figure 3 with some exceptions as follows. **(a)**: Blue circles denote households with *ω*_overall_ < 1 with the size of the circles showing the magnitude of 1/(5 × *ω*_overall_) to make their sizes comparable to the pink circles. Households with the largest *ω*_overall_ include HH23, HH18, and HH58. **(d)** A moderate seasonal signal was observed in Kanungu.

In Tororo, a seasonal varying hotspot was identified for the dry vs. rainy seasons comparison while six households were identified as seasonal varying hotspots before the deployment of IRS vs. after IRS (Figure 5(b-c)). Tororo, the study site with highest transmission intensity, exhibited a strong seasonal signal in mosquito counts over time (Figure 5(d)). Here we see clearly that mosquito counts peaked at the end of rainy season and at the very beginning of dry season and reduced remarkably after the deployment of IRS. Most households in Tororo behaved very similarly during dry and rainy seasons but showed large differences in *ω* before the deployment of IRS and after (Figure 5(e-f)).

**Figure 5.**
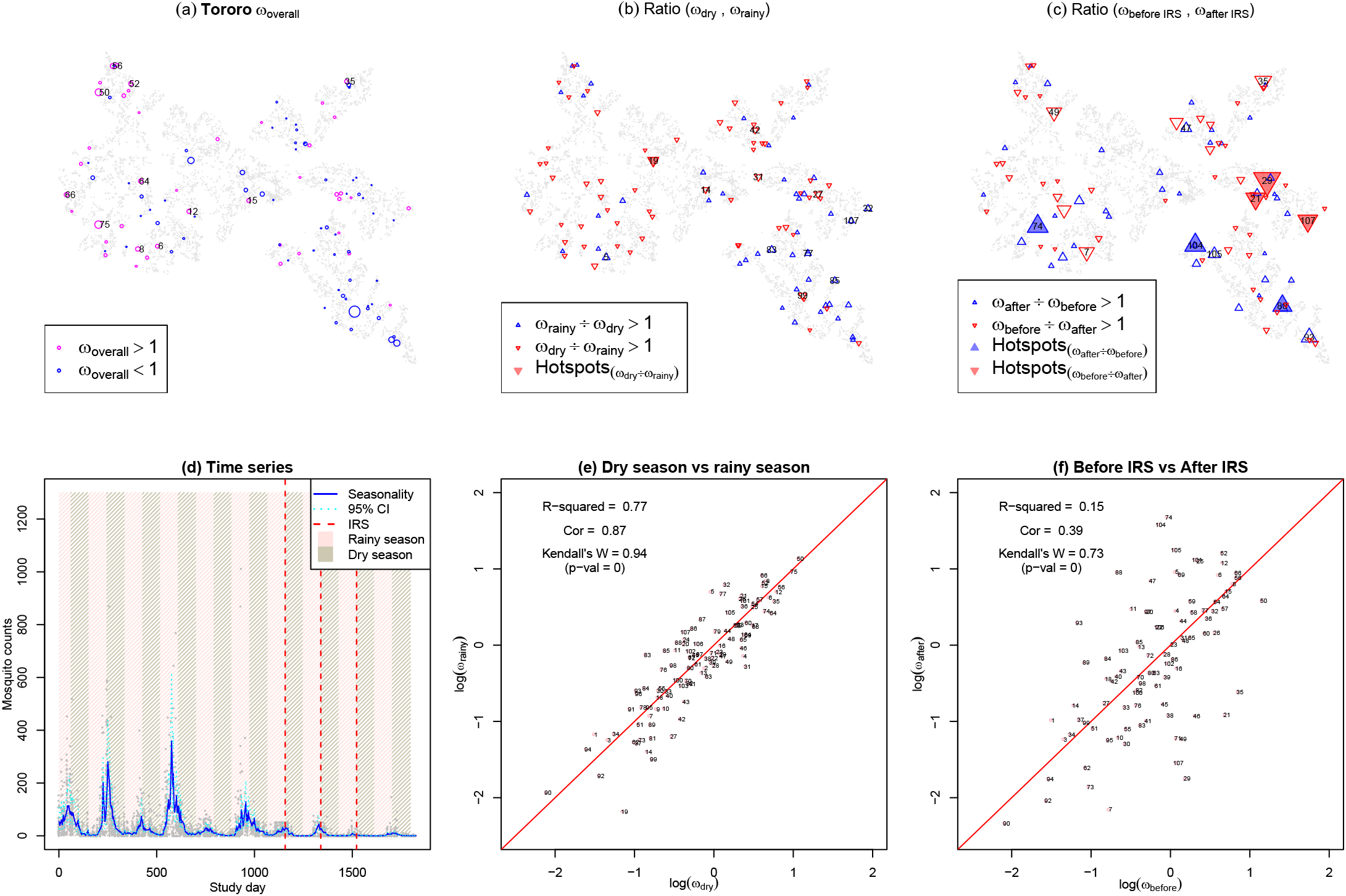
Tororo. The layout of the figure is the same as in Figure 3 with some exceptions as follows. **(a)**: Blue circles denote households with *ω*_overall_ < 1 with the size of the circles showing the magnitude of 1/(2× *ω*_overail_) to make their sizes comparable to the pink circles. Households with the largest *ω*_overail_ include HH75, HH50, HH56, and HH52. **(c)** Ratios of household biting propensities before the deployment of IRS (*ω*_before KS_) and after the deployment of IRS (ω_after IRS_). Each plotted upward pointing blue triangle represents a household with a larger *ω* after the deployment of IRS compared to before the deployment of IRS; a blue filled triangle denotes a hotspot for the ratio of *ω*_after IRS_/*ω*_beforeIRS_· Each plotted downward pointing red triangle represents a household with a larger *ω* before the deployment of IRS compared to after the deployment of IRS; a red filled triangle denotes a hotspot for the ratio of *ω*_beforeIRS_/*ω*_after IRS_· **(d)** The red vertical dashed lines denote the deployment of IRS which were six months apart. A strong seasonal signal was observed in Tororo. **(f)** Scatter plot of log(ω_beforeIRS_) and log(ω_after IRS_) along with measures of *R*^2^, correlation, and Kendall’s W.

### Association of heterogeneous household biting with environmental and housing covariates

High resolution climatic and environmental covariates (100m × 100m) known to interact with mosquito density, along with housing covariates for all the sampled households at the three study sites were assembled (Table 1). For each of the environmental covariates, we calculated and plotted Pearson correlation coefficients for the covariate and *ω* of various scenarios to understand the association of a covariate on mosquito biting propensities (Figure 6). The environmental covariates did not have strong associations with household biting heterogeneity at all three sites. Nevertheless, the households in Kanungu had greater associations of biting propensities with environmental covariates including land surface temperature (day time, night time, and diurnal difference), precipitation, and elevation, compared to two other sites. Kanungu was a rural area of rolling hills in western Uganda located at an elevation of 1,310 m above sea level and it was the study site with the highest elevation and greatest within-site elevation change among the three sites (Maxwell et al., 2014). The greater variation in environmental characteristics across households in Kanungu due to its altitude gradient most likely contributed to the greater association between biting propensities and the environmental covariates.

**Figure 6.**
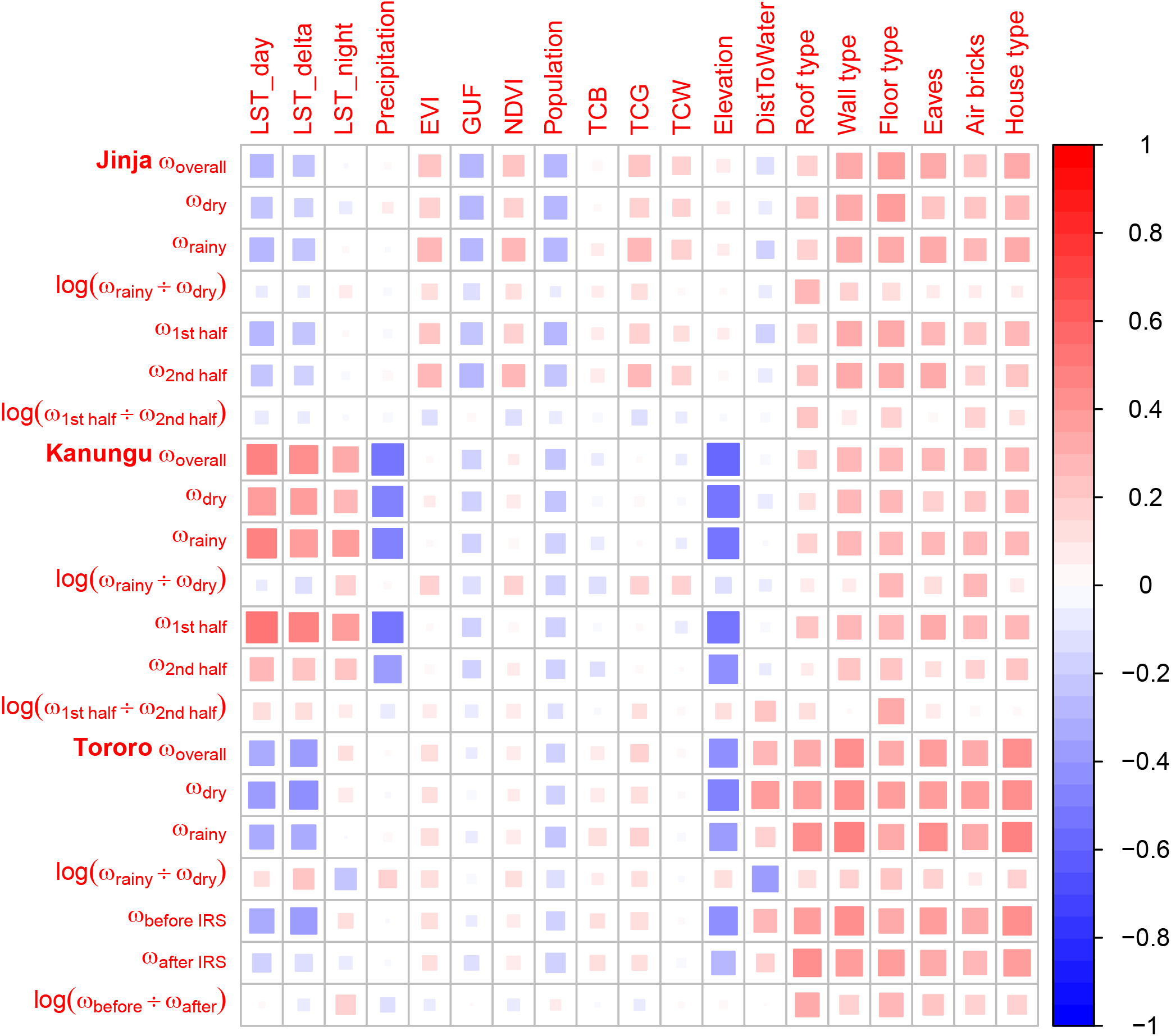
Correlation heatmap of household biting propensities and environmental and housing covariates. See Table 1 for descriptions of the covariates. The size of the squares and the shade of colors both illustrate the magnitude of the correlation. Red-shaded squares denote positive correlation whereas blue-shaded squares denote negative correlation.

**Table 1.**
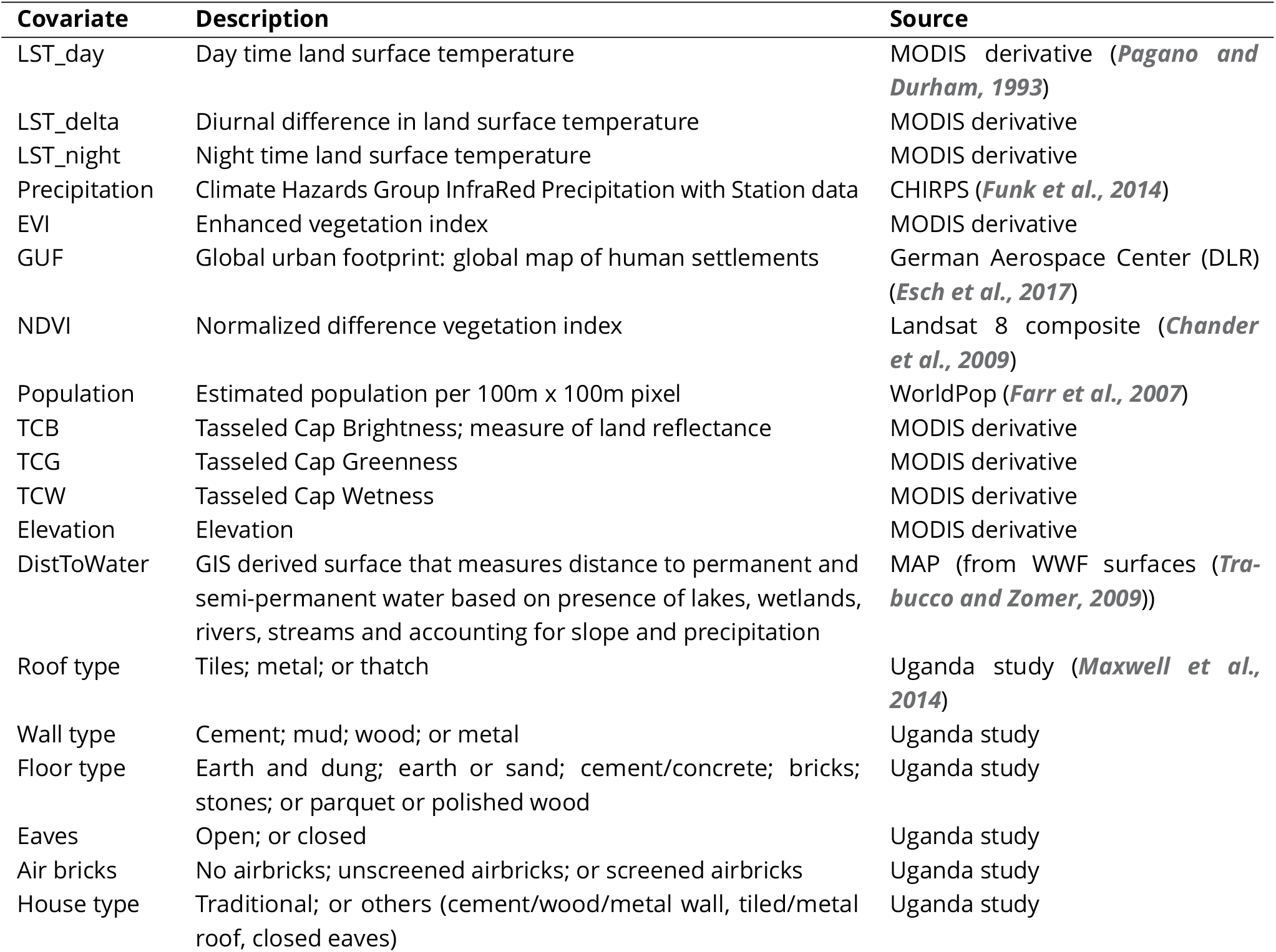
Description of environmental and housing covariates.

The housing covariates (roof type, wall type, floor type, eaves, air bricks, and house type) were all available as categorical variables, we therefore fitted a linear regression model for the housing covariates and *ω* of various scenarios, which resulted in multiple correlation coefficient for each pair. Among the three sites, the housing covariates had the strongest associations with *ω* in Tororo, followed by *ω* in Jinja, and *ω* in Kanungu. Maps of Tororo in Figure S3 show the average predicted *Anopheles* mosquito (*An. gambiae (s.l.)* and *An. funestus (s.l.)*) density per household per night across the whole study region before the deployment of IRS and after the deployment of IRS. In separate analyses, the predictions were obtained by fitting counts of malaria vectors using a zero-inflated negative binomial model in a Bayesian geostatistical spatio-temporal framework along with environmental covariates (Alegana et al., 2016). Clearly, the environmental covariates did not account for a big portion of the heterogeneity in mosquito density, especially in the density of *An. gambiae (s.l.)* before the deployment of IRS (upper left panel of Figure S3). A big part of the heterogeneity in mosquito biting was attributed to household attractiveness caused by housing characteristics or structures.

For each study site and for all three study sites combined, a separate negative binomial regression was used to model the relationship between the six categorical housing covariates and the total number of *Anopheles* mosquito caught per household by light trap catches, with the number of sampling nights included as an offset term in the model. Table 2 summarizes the incidence rate ratios (IRR) of the best fitted model (retaining significant risk factors only) selected by the Akaike information criterion. In Jinja, houses with mud, wood, and metal walls had a higher malaria risk compared to houses with cement walls; houses with earth or sand floors in Jinja were also much more likely to attract mosquitoes compared to other types of floors. In Kanungu and Tororo, houses with mud floors had a higher malaria risk compared to houses with cement floors; houses with open eaves also attracted a lot more mosquitoes compared to houses with closed eaves. On the other hand, houses with screened airbricks in Tororo had a protective effect towards mosquito biting or malaria risk. For all sites combined, roof type was a highly significant risk factor in addition to wall type, floor type, and airbricks. Houses with thatched roofs were a lot more likely to attract mosquitoes compared to houses with metal or tiled roofs.

**Table 2.** Association between household characteristics and the anophelines collected per household per night (total anophelines caught / total nights of collection) at three sites in Uganda, estimated with negative binomial regression models. The best fitted model had identified four risk factors of household biting for all sites combined (wall type, floor type, airbricks, and roof type); two risk factors for Jinja (wall type and floor type); two risk factors for Kanungu (wall type and eaves); and three risk factors in Tororo (wall type, eaves, and airbricks). (IRR: incidence rate ratio; CI: confidence interval)

|  |  | All sites combined |  | Jinja |  | Kanungu |  | Tororo |  |
| --- | --- | --- | --- | --- | --- | --- | --- | --- | --- |
| Characteristics |  | IRR (95% CI) | p | IRR (95% CI) | p | IRR (95% CI) | p | IRR (95% CI) | p |
| Wall type | Cement | 1.00 | - | 1 | - | 1 | - | 1 | - |
|  | Mud | 1.70 (0.99-2.92) | 0.069 | 1.69 (0.80-3.58) | 0.187 | 1.99 (1.16-3.36) | 0.010 | 1.96 (1.32-2.93) | <0.001 |
|  | Wood | 0.18 (0.07-0.58) | 0.001 | 2.27 (0.92-6.31) | 0.067 | - | - | - | - |
|  | Metal | 0.60 (0.20-2.31) | 0.377 | 4.23 (1.65-12.42) | 0.003 | - | - | - | - |
| Floor type | Earth or sand | 1 | - | 1 | - |  |  |  |  |
|  | Earth and dung | 2.23 (1.49-3.39) | <0.001 | 0.94 (0.47-2.00) | 0.865 |  |  |  |  |
|  | Parquet or polished wood | 0.37 (0.04-44.68) | 0.509 | 0.19 (0.03-4.80) | 0.160 |  |  |  |  |
|  | Bricks | 0.07 (0.01-1.28) | 0.013 | - | - |  |  |  |  |
|  | Cement/concrete | 0.74 (0.45-1.24) | 0.311 | 0.42 (0.20-0.86) | 0.022 |  |  |  |  |
|  | Stones | 0.46 (0.09-7.73) | 0.443 | - | - |  |  |  |  |
| Eaves | Open | - | - |  |  | 1 | - | 1 | - |
|  | Closed | - | - |  |  | 0.56 (0.32-0.96) | 0.038 | 0.74 (0.58-0.97) | 0.026 |
| Airbricks | None | 1.00 | - |  |  |  |  | 1 | - |
|  | Unscreened | 1.12 (0.71-1.75) | 0.592 |  |  |  |  | 1.17 (0.82-1.68) | 0.355 |
|  | Screened | 0.18 (0.09-0.42) | <0.001 |  |  |  |  | 0.30 (0.13-0.79) | 0.006 |
| Roof type | Thatch | 1.00 | - |  |  |  |  |  |  |
|  | Metal or tiles | 0.44 (0.27-0.70) | 0.001 |  |  |  |  |  |  |

## Discussion

We have shown that it is possible to quantify heterogeneity in malaria exposure and its component parts: seasonality and household biting propensities. The proposed method was successfully validated via an extensive simulation study. Household biting propensities, seasonality, and environmental factors could be quantified using a Bayesian ZINB regression model along with a temporal smoothing prior distribution for estimating seasonal signal. Using these biting propensities, we were able to identify the factors accounting for differences in mosquito counts at three study sites in Uganda. Despite the variance introduced by seasonality, biting propensities tended to be strongly correlated with the average counts within a site, though there were some notable outliers.

While many studies of malaria transmission have focused on quantifying spatial patterns or seasonality, substantially less attention has been paid to quantifying heterogeneous biting propensities among individuals or households or environmental noise and its importance for malaria transmission. While some evidence suggests biting patterns are spatially heterogeneous due to environmental factors (Takken and Lindsay, 2003), these patterns had not been examined critically alongside other measures of heterogeneity, such as seasonality, household quality or other factors, and environmental noise. Our study, like others (Bejon et al., 2014), has shown that heterogeneity is operating at multiple scales. Large differences among households, which are partly explained by household quality, are also influenced by spatial patterns related to other environmental covariates (e.g. elevation (altitude gradient) and urbanicity), which are likely related to mosquito ecology. Our method quantifies heterogeneity at a small scale (i.e. at the household level), validated by simulation studies, giving us confidence in our analysis of complex spatial patterns collected in field studies.

Our work provided an improved understanding of heterogeneity in malaria exposure at the three study sites in Uganda and offered a valuable opportunity for assessing whether interventions could be spatially targeted to be aimed at households with the highest mosquito biting propensities or spatial areas, i.e., hotspots of malaria risk. While analysis using scan statistics can identify statistical anomalies, our method is better at quantifying pattern and identifying its causes. We are able to identify cold spots (households with very small biting propensities) as well as hot spots (households with very large biting propensities). The study showed that the presence of malaria hotspots was associated with the environment (including urbanicity, distance to water bodies or aquatic habitats, rainy seasons, and elevation) as well as housing characteristics. It has been found that house design in rural Uganda was associated with additional reductions in mosquito density and parasite prevalence following the introduction of IRS (Rek et al., 2018). In the Ugandan context, malaria control efforts should be targeted towards houses with features that led to an elevated malaria risk, including houses with mud walls, earth or sand floors, earth and dung floors, open eaves, unscreened airbricks, and thatched roofs. Improvements on housing design such as house-screening, closing the eaves, or installing ceilings help prevent mosquitoes gaining access to houses and should therefore reduce malaria transmission and infection (Njie et al., 2009; Lindsay et al., 2002).

Indeed, assessing the likely benefits of spatial targeting requires some quantitative understanding of heterogeneity and its underlying causes, including seasonality, factors that make some households more attractive or easier to enter than others, environmental noise, and measurement error. Focusing malaria control efforts on households that contribute disproportionately to malaria transmission could help achieve community protection by eliminating transmission in a relatively small fraction of human hosts (Woolhouse et al., 1997; Smith et al., 2007). If heterogeneous transmission is assumed and high-risk households were missed, control efforts would be relatively inefficient (Bousema et al., 2012). Evidence-based targeting of malaria control interventions on high-risk individuals and hotspots could bring a wide range of economic benefits for the countries concerned. Combining malaria control interventions such as indoor spraying with residual insecticides, artemisinin-based combination therapy, indoor residual spraying, and long lasting insecticidal bednets are known to be an effective way for reducing malaria burden, for instance, in Zanzibar (Bhattarai et al., 2007), Western Kenya (Stuckey et al., 2014), and at Tororo in Uganda (Katureebe et al., 2016).

Differences in vector seasonality were apparent at the three study sites. Based on the results of our study, Tororo (the most rural site) had the strongest seasonality, while Jinja (where mosquitoes were sampled in the town) had the weakest seasonality. The magnitude of seasonality at the three sites appeared to correlate with mosquito densities over time. The dynamics and seasonal abundance of malaria vectors are known to be associated with micro-ecology, rainfall and temperature patterns (Dery et al., 2010). Surface water and temperature are among the two major drivers that affect mosquito abundance, blood feeding rates, and parasite development within vectors (Mordecai et al., 2013). While seasonal patterns are a common feature of mosquito abundance and malaria transmission, weather is the proximal cause and it is as likely to cause inter-annual deviations from the canonical seasonal pattern (Weiss et al., 2014). The increased availability of aquatic habitats during rainy seasons often results in high vector abundance, but on the other hand, it can also lead to flooding and cause the aquatic stages of mosquitoes to be washed out (Bisanzio et al., 2015). The presence of seasonal or permanent aquatic habitats such as paddies or ponds and degree of urbanization of a site also contributes to differences in mosquito abundance across geographies (Parham and Michael, 2010).

The households in Kanungu and Tororo mostly had similar biting propensities during different seasons while households in Jinja showed greater differences in *ω* during different seasons. Cleary, the landcover in Jinja town would not support the temporary breeding sites that follow rainy seasons in rural areas. The identification of three hotspots in Jinja compared to only one hotspot each in Kanungu and Tororo for the dry vs. rainy seasons comparison suggested that mosquito biting attractiveness of households in Jinja might be more susceptible to changes in seasons than in Kanungu and Tororo. This could possibly be attributed to the fact that the sampled households in Jinja were in close proximity to a swampy area near Lake Victoria, which acted as a critical water body (see Figure S4). Distance of houses to water bodies had long been known as a risk factor of malaria (Kleinschmidt et al., 2001). These findings suggested that it would be possible to target different households during different seasons for optimal control efforts.

Household mosquito counts recorded in Jinja in the second half period of surveillance (1,172 mosquitoes) were only about half of those recorded in the first half period (2,309 mosquitoes). Similarly, household mosquito counts recorded in Kanungu in the second half period (5,177 mosquitoes) were only about half of those recorded in the first half period (10,018 mosquitoes). Such observations suggested some changes in malaria transmission over time at both study sites, however, Katureebe et al. (2016) found no association of LLIN with malaria transmission in Jinja and only modest effect of LLIN in Kanungu. A remarkable reduction of mosquito abundance after the deployment of IRS was observed in Nagongera where IRS began in December 2014 (see Figures S2 and S3), which at the same time reduced mosquito biting propensities in about two-third of the households. There were three hotspots with a much larger *ω* before IRS compared to after IRS (HH21, HH29, and HH107); these households were located reasonably close to each other, i.e. in the middle right area of the study site. The three hotspots with a much larger *ω* after IRS compared to before IRS were HH74, HH88, and HH104 which were located across the lower area of the study site. Despite a drastic reduction in household mosquito counts after IRS in these three households, i.e. from 1,295 mosquitoes to 223 mosquitoes in HH74, from 999 mosquitoes to 109 mosquitoes in HH88, and from 1,211 mosquitoes to 177 mosquitoes in HH104, biting propensities in these households after IRS increased instead of decreased, relative to other households.

In conclusion, the study found that housing quality contributed to a large portion of the heterogeneity in household mosquito biting. Cement walls, brick floors, closed eaves, screened airbricks, and tiled roofs are features of a house that had shown protective effects towards malaria risk. Household mosquito biting propensities in Jinja showed some important differences during the dry season and the rainy season, most likely due to the close proximity of the study site to a swampy area. Jinja and Kanungu had shown lower malaria transmission in the last half of the study, supported by the reduction of mosquito densities in the second half period of surveillance. The application of IRS in Tororo had caused a massive reduction in mosquito abundance as well as reducing household biting propensities in two-third of the households. Based on these findings, IRS was a successful measure of vector control interventions in Uganda and improvements in house quality should be recommended as a supplementary measure for malaria control.

## Materials and methods

### Data sources

Entomological surveillance was conducted in three areas of Uganda: between October 2011 and March 2015 for Walukuba subcounty, Jinja District (0°26’33.2” N, 33°13’32.3” E)and Kihihi subcounty, Kanungu District (0°45’3.1” S, 29°42’3.6” E); and between October 2011 and September 2016 for Nagongera subcounty, Tororo District (0°46’ 10.6” N, 34°1’34.1” E). Details of this study, including the overall study design, study sites, a detailed description of the entomological methods, and ethical approval have been described elsewhere (Maxwell et al., 2014; Kamya et al., 2015). See Figures S4 to S6 for the maps of the three study sites.

Jinja District is located in the east central region bordering Lake Victoria and has a population of roughly 470,000 with 63% of the population living in rural area (Uganda Bureau of Statistics, 2016). Jinja town was characterized historically by moderate malaria transmission intensity, but it was the site with lowest transmission in our study. Kanungu District is a rural area in south-western Uganda bordering the Democratic Republic of Congo and has a population of approximately 250,000 with 80% of the population living in villages (Uganda Bureau of Statistics, 2016). Malaria in Kanungu has been characterized by relatively low transmission intensity and low endemicity, but it had moderate transmission in our study. Tororo District is located in south-eastern Uganda on the Kenyan border and has a population of about 520,000 with 86% of the population living in villages (Uganda Bureau of Statistics, 2016). Tororo is characterized by very high malaria transmission.

CDC light traps (Model 512; John W. Hock Company, Gainesville, FL, USA) were placed in 330 randomly-selected households; 116 in Jinja, 107 in Kanungu, and 107 in Tororo. The traps were positioned one meter above the floor at the foot of the bed where a study participant slept. Traps were set on one day each month at 19:00 h and collected the following morning at 07:00 h. All anophelines were identified taxonomically based on morphological criteria according to established taxonomic keys (Gillies and Coetzee, 1987). Up to 50 anophelines per household were tested for presence of *Plasmodium falciparum* sporozoites using ELISA (Wirtz et al., 1989). The total number of anophelines trapped in Jinja per household each day ranged from 0 to 121 with a median of 0; ranged from 0 to 306 with a median of 0 in Kanungu; and ranged from 0 to 1011 with a median of 9 in Tororo.

### Statistical analyses

Here we partitioned the observed variance of mosquito counts at the three Ugandan study sites among the sources of heterogeneity attributed to individual households, seasonality, and environmental noise or measurement error. Here, ‘partitioning’ refers to attributing proportions of the total variability to individual factors. Partitioning the total variance of a response variable into its component sources is common in ecological studies (Fletcher and Underwood, 2002; Qian and Shen, 2007) as it can help inform a wide variety of research and management questions (Lindenmayer and Likens, 2009; Irwin et al., 2013). For instance, the variance partitioning approach developed by Irwin et al. (2013) for a negative binomial mixed model was a useful method for assessing the response of a variance structure to large-scale ecological changes. On the other hand, as observed in our mosquito data, overdispersion is commonly exhibited in ecological count data, and can be modeled effectively using a variety of methods, including a negative binomial distribution.

The mosquito counts consisted of a large proportion of zero counts; 70% in Jinja, 54% in Kanungu, and 21% in Tororo; and showed some degree of overdispersion. As a general rule, the presence of over 30% of zeros in the data would require a zero-inflated model. We adopted a probability model that is capable of handling excess zeros while modeling non-zero counts properly, namely a zero-inflated negative binomial model (ZINB) (Lambert, 1992). A Bayesian framework was adopted to estimate model parameters, using the integrated nested Laplace approximation approach (Rue et al., 2009). By fitting the ZINB regression models to the mosquito counts, we estimated household biting propensities (*ω*), seasonal signal (*S*), and noise (*e*) at each of the study sites.

For mosquito counts ***Y*** = {*y*_1_, *y*_2_,…, *y*_n_, the ZINB distribution can be written as

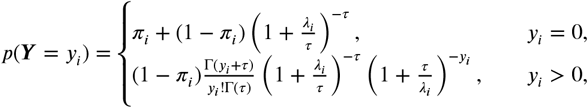

where τ > 0 is a shape parameter which quantifies the amount of overdispersion. The mean and variance of the ZINB distribution are *E*(*Y*_i_) = (1 - *π*_i_)*λ*_i_ and var(*Y*_i_) = *λ*_i_(1 - *π*_i_)(1 + *π*_i_*λ*_i_ + *λ*_i_/*τ*), respectively.

For each of the study sites, the expected count for household *j* on day *i, λ*_ij_, was modeled on a set of explanatory variables with the aid of a log link function, while the probability *π*_ij_ was modeled on the same set of explanatory variables using a logit link function. The covariates of interest included household identifiers (ID) and sampling days (*t*). Household identifiers were treated as fixed effects in the regression model, whereas daily seasonal signal and random noise were treated as random effects. A range of Bayesian prior distributions were imposed on sampling days (*t*), including first- and second-order random walks (RW1, RW2) and autoregressive processes of order 1 and order 2 (AR1, AR2) (Sørbye and Rue, 2014). The random noise was assumed to be independent and identically distributed (i.i.d.). The Watanabe-Akaike information criterion (Watanabe, 2013) was used as the model comparison criterion in order to select the model that produced the best estimates of *ω, S*, and *e* for each of the three study sites in Uganda. Note that the estimated *ω* for each of the following scenarios had been scaled to have a mean of one so that they reflected relative attractiveness of household mosquito biting with respect to other households.

The model for estimating the overall household biting propensities (*ω*_overall_) is as follows,

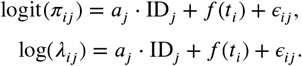

Here *a_j_* quantifies the effects of household biting. The random noise, *Є*_ij_, are assumed to be i.i.d., and *t_i_*, the temporally structured random effects, are assigned one of the four Bayesian temporal smoothing prior distributions. Using the estimated parameters, we obtained the household biting propensities, *ω*_overall_ = exp(*a_j_*) and the seasonal signal, *S* = exp(/*f*(*t_i_*)). The noise, *e* = exp(*Є_ij_*), accounted for additional variation among mosquito counts that was not accounted for by the covariates or by Poisson (random) variation around the mean *ωS*.

Household biting propensities during dry (*ω*_dry_) and rainy (*ω*_rainy_) seasons, during the first half period of surveillance (*ω*_1st half_) and the second half period (*ω*_2nd half_), and before the deployment of IRS (*ω*_beforeIRS_) and after the deployment of IRS (*ω*_after IRS_) were estimated using one of the following sets of equations,

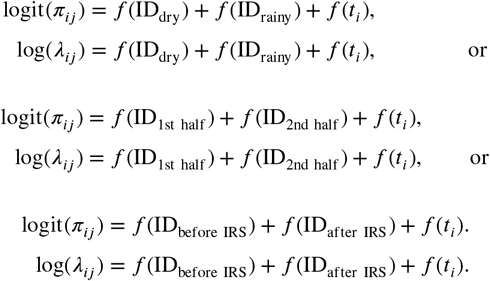

Here, ID_dry_ denote household IDs during dry season, ID_rainy_ denote household IDs during rainy season, and so forth. The random effects *f*(ID_…_) was assumed to be i.i.d. By modeling the biting propensities of different scenarios as random effects, it allowed for estimation and comparison of two scenarios within one model, instead of having to split the datasets into two subsets. The noise term, Є_ij_, was not required in the model because the focus here was to estimate household-level effects. Hence, the biting propensities for different scenarios are such that: *ω*_dry_ = exp(*f*(ID_dry_)); *ω*_rainy_ = exp(*f*(ID_rainy_)); *ω*_1st half_ = exp(*f*(ID_1st half_)); *ω*_2nd half_ = exp(*f*(ID_2nd half_)); *ω*_before IRS_ = exp(*f*(ID_before IRS_)); and *ω*_after IRS_ = exp(*f*(ID_safter IRS_)).

We also conducted hotspot analysis on *ω* using the Getis-Ord (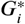) statistic (Getis and Ord, 1992) for all three sites to identify hotspots of malaria risk. The 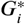 statistic is a local statistic that identifies variation across the study area, by focusing on individual features and their relationships to nearby features. It also identifies statistically significant hot spots and cold spots, given a set of weighted features. To be statistically significant, the hot spot or cold spot will have a high or low value and be surrounded by other features with high or low values. The value of the target feature is included in analysis. The 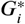 statistic is a z-score and given as:

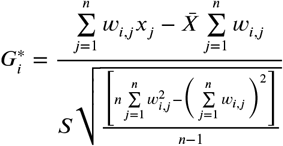

where *x_j_* is the attribute value for feature *j*(*x_j_* = *ω_j_* in this case), *w_i,j_* is the spatial weight between feature *i* and *j, n* is the total number of features while:

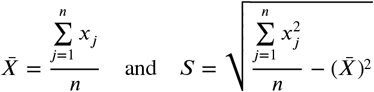

The Getis-Ord teslas performed using the localG function in the spdep package in R (Bivand et al., 2011). At a significance level of 0.001, a z-score would have to be less than −3.29 or greater than 3.29 to be statistically significant.

## Additional information

### Supplementary files

Supplementary File 1: Supplementary figures.

Supplementary File 2: Detailed description of simulation analyses for identifying the most robust method for quantifying heterogeneity in malaria exposure.

### Funding

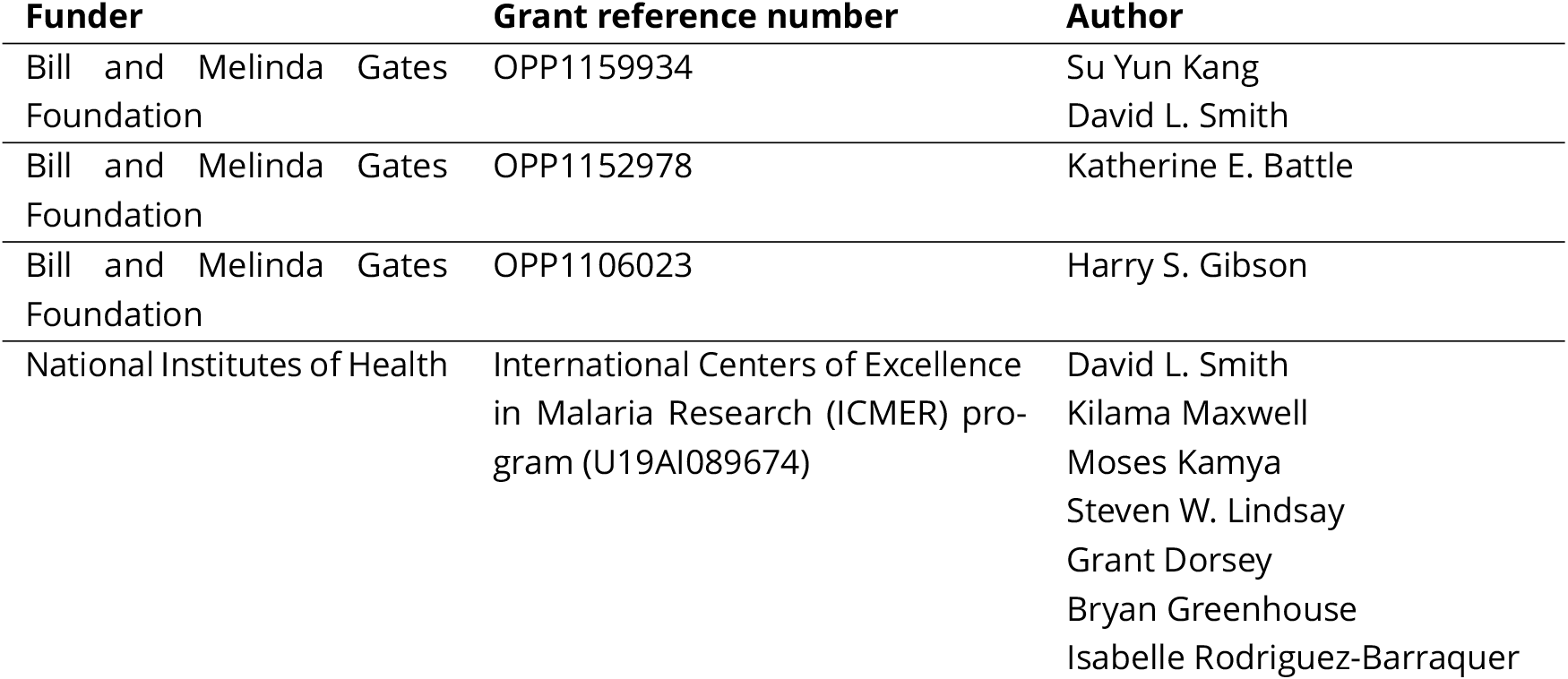

### Author contributions

SYK, DLS, DB, Conception and design, Analysis and interpretation of data, Drafting or revising the article; KEB, HSG, Acquisition of data, Drafting or revising the article; LVC, KM, MK, SWL, GD, BG, IRB, RCR, Drafting or revising the article;

## Acknowledgments

The authors would like to thank those in the study communities who participated in this study or helped collect mosquitoes. The authors wish to acknowledge the Infectious Diseases Research Collaboration (IDRC) for administrative and technical support.

## Supplementary File 1: Supplementary Figures

**Kang *et al***.

**Figure S1.**
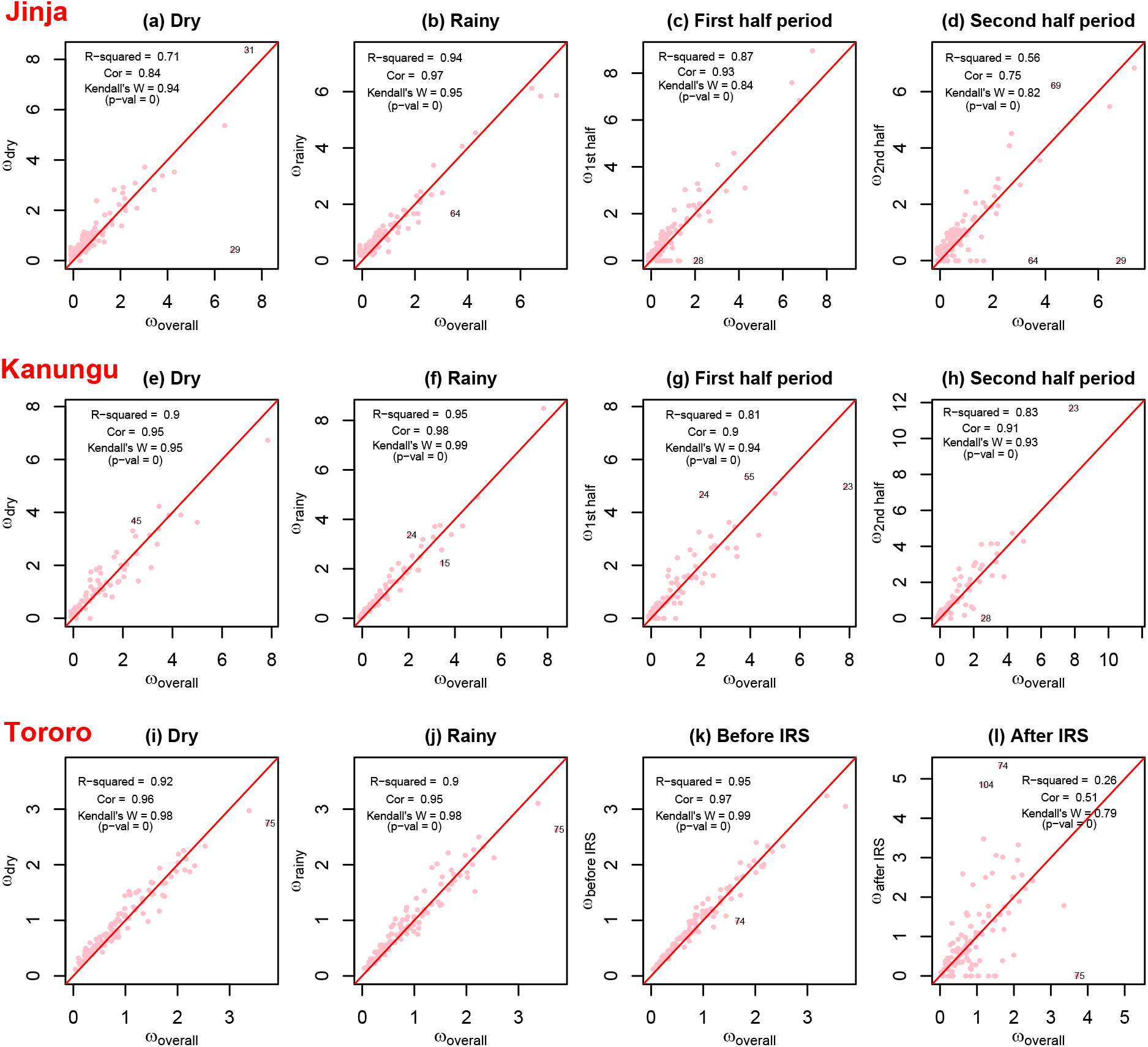
Household biting propensities for different scenarios are plotted against the overall biting propensities (*ω*_overall_) for Jinja, Kanungu, and Tororo (top to bottom rows), along with measures of ***R***^2^, correlation, and Kendall’s *W*. From left to right columns: *ω*_overall_ against biting propensities during dry season (*ω*_dry_); *ω*_overall_ against biting propensities during rainy season (*ω*_rainy_); *ω*_overall_ against biting propensities during the first half period of surveillance (*ω*_1st half_) or before the enrollment of IRS (*ω*_before ms_); and *ω*_overall_ against biting propensities during the second half period of surveillance (*ω*_2rd half_) or after the enrollment of IRS (*ω*_after IRS_). We fitted a linear model for *ω*_overall_ and ***ω*** observed for different scenarios and used the `outlierTest` function in the `car` library in `R` to identify outliers given the fitted model. The most extreme observations based on the given model were labelled with the household (HH) number. Among all the outlier households, the most notable ones that appeared in more than one scenario were HH29 and HH64 in Jinja; HH23 and HH24 in Kanungu; and HH74 and HH75 in Tororo.

**Figure S2.**
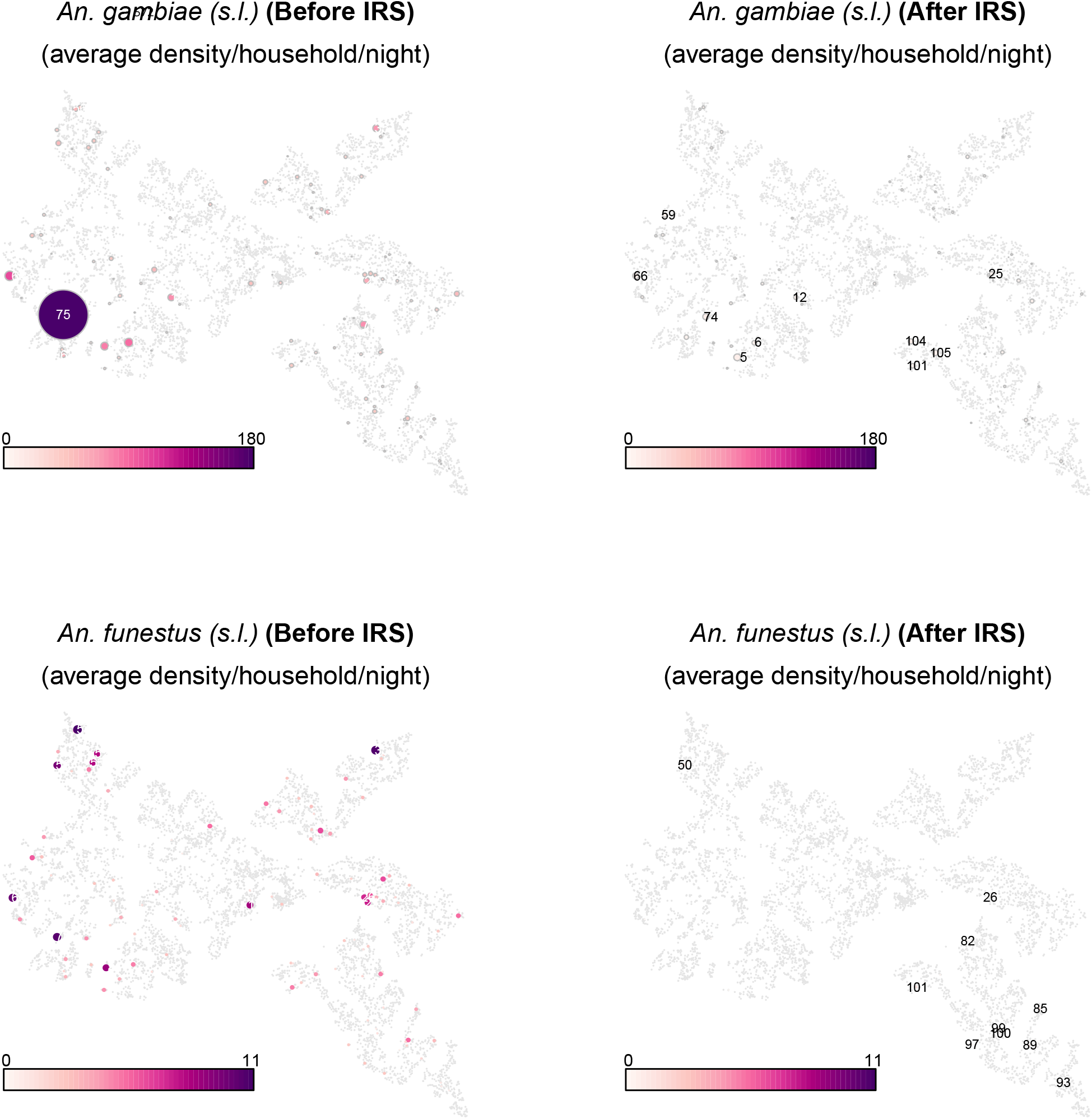
Tororo. *Anopheles* mosquitoes were made up of too main malaria vector species: *An. gambiae* (*sensu lato*) and *An. funestus* (*sensu lato*). The dots on each map show the average *An. gambiae* (*s.l.*) and *An. funestus* (*s.l.*) density per household per night before the enrollment of IRS and after the enrollment of IRS. *An. gambiae* (*s.l.*) is a dominant malaria vector species in Tororo. There was a remarkable reduction of mosquito abundance after the enrollment of IRS for both vector species. Households with the highest mosquito density were labelled with the household number.

**Figure S3.**
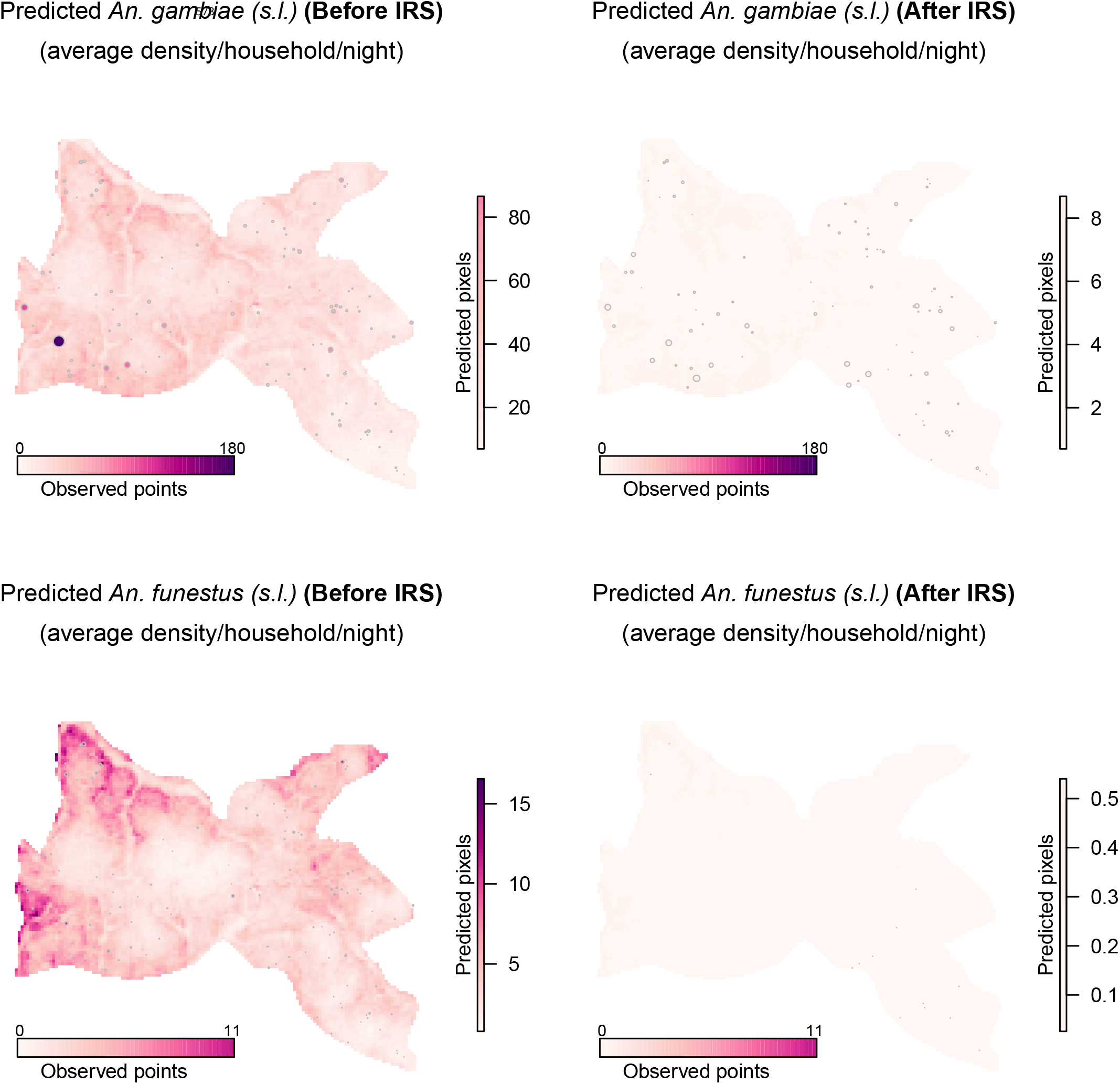
Tororo. The maps show the average predicted *An. gambiae* (*s.l.*) and *An. funestus* (*s.l.*) density per household per night across the whole study region before the enrollment of IRS and after the enrollment of IRS. The predictions were obtained by fitting counts of malaria vectors using a zero-inflated negative binomial model in a Bayesian geostatistical spatio-temporal framework along with environmental covariates. The circles overlaid on the maps denote the average observed density of malaria vectors per household per night. The color legend on the right of each panel represents the average predicted density while the color legend at the bottom of each panel represents the average observed density.

**Figure S4.**
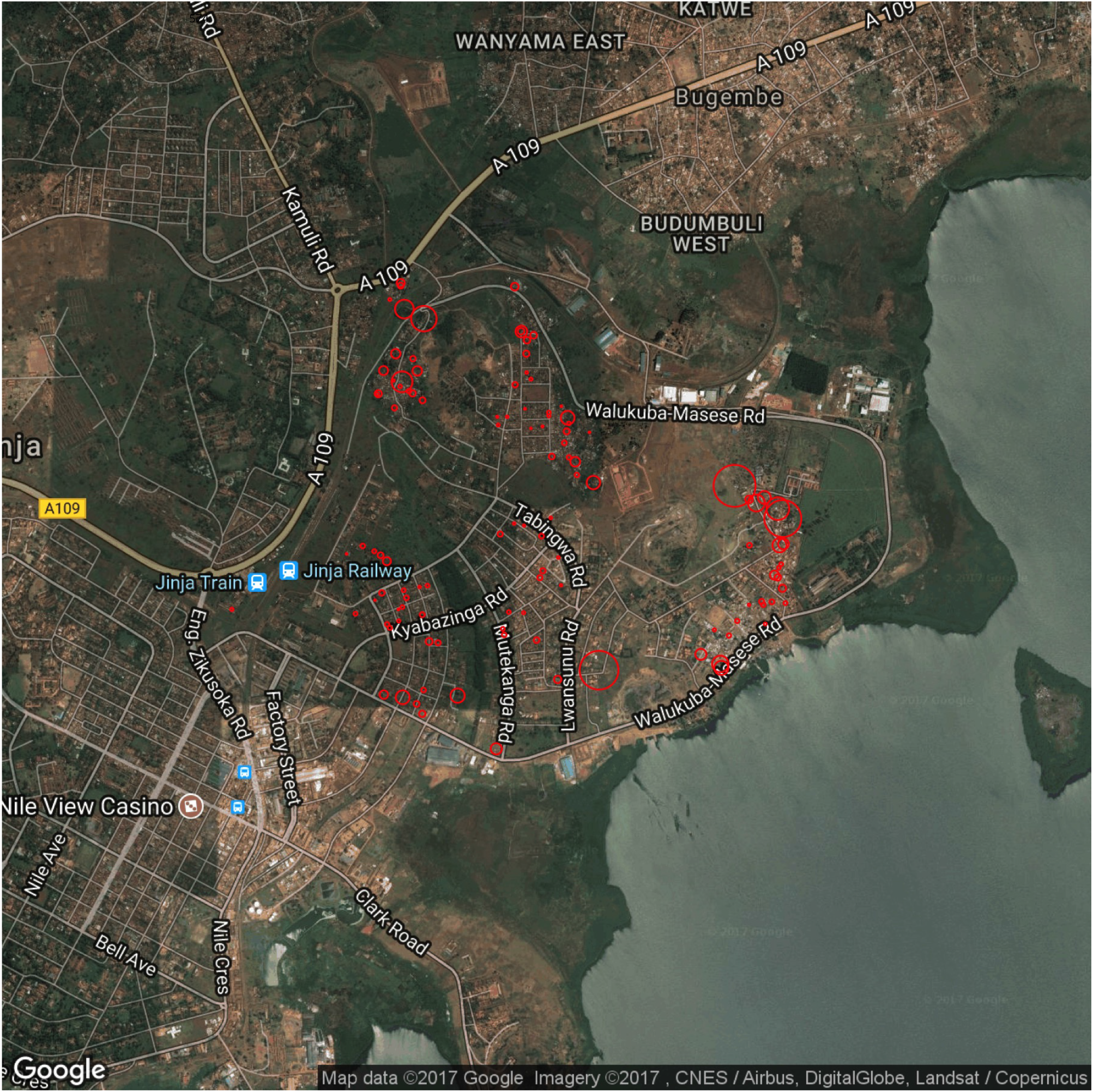
Sampled households in Walukuba subcounty,Jinja District between October 2011 and March 2015, located in the east central region bordering Lake Victoria. Each red circle denotes a household and the size ofthe circles denotes the overall biting propensities over the entire duration of surveillance. This is the site with lowest malaria transmission in our study.

**Figure S5.**
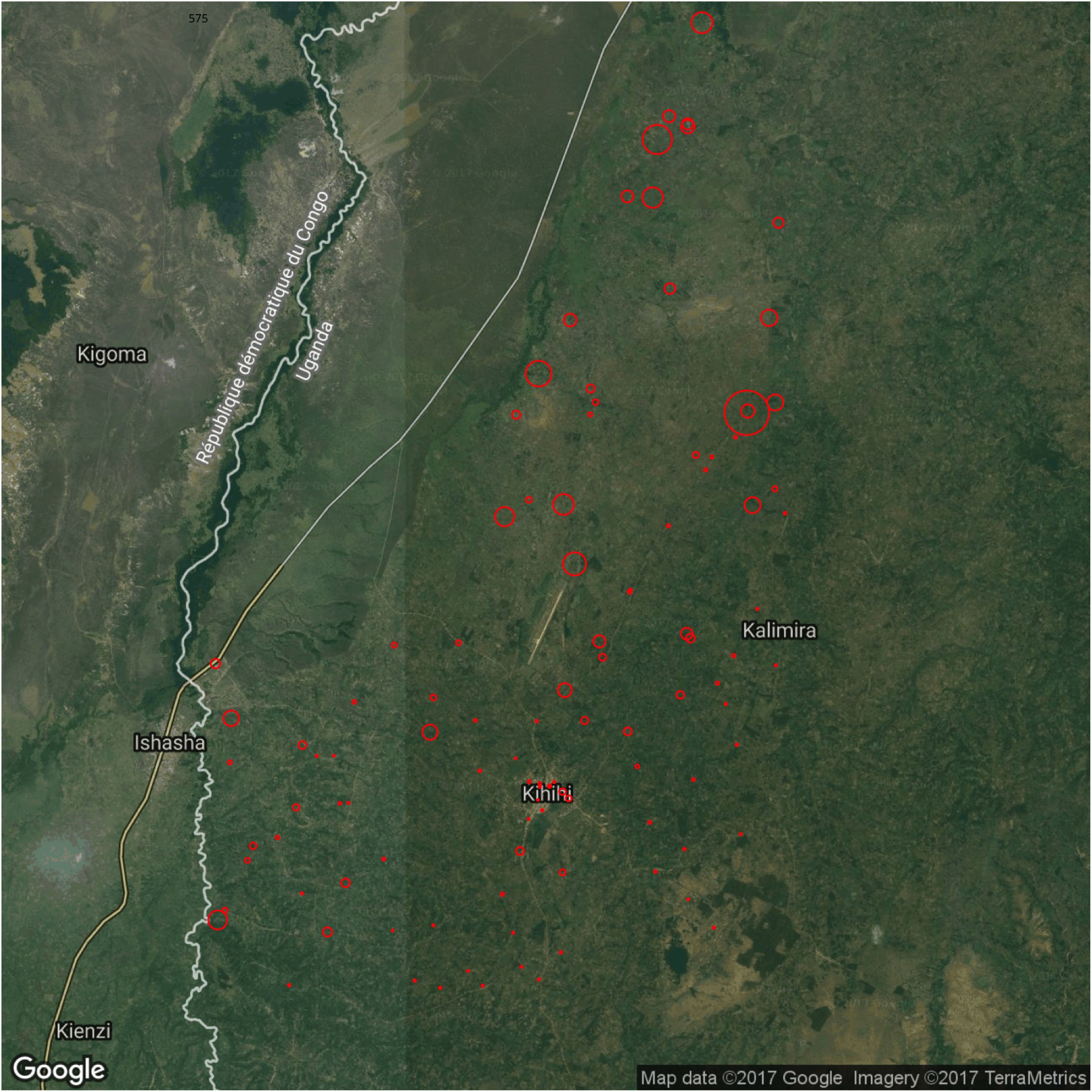
Sampled households in Kihihi subcounty, Kanungu District between October 2011 and March 2015, located in a rural area in south-western Uganda bordering the Democratic Republic of Congo. Each red circle denotes a household and the size ofthe circles denotes the overall biting propensities overthe entire duration ofsurveillance. This is the site with moderate malaria transmission in our study.

**Figure S6.**
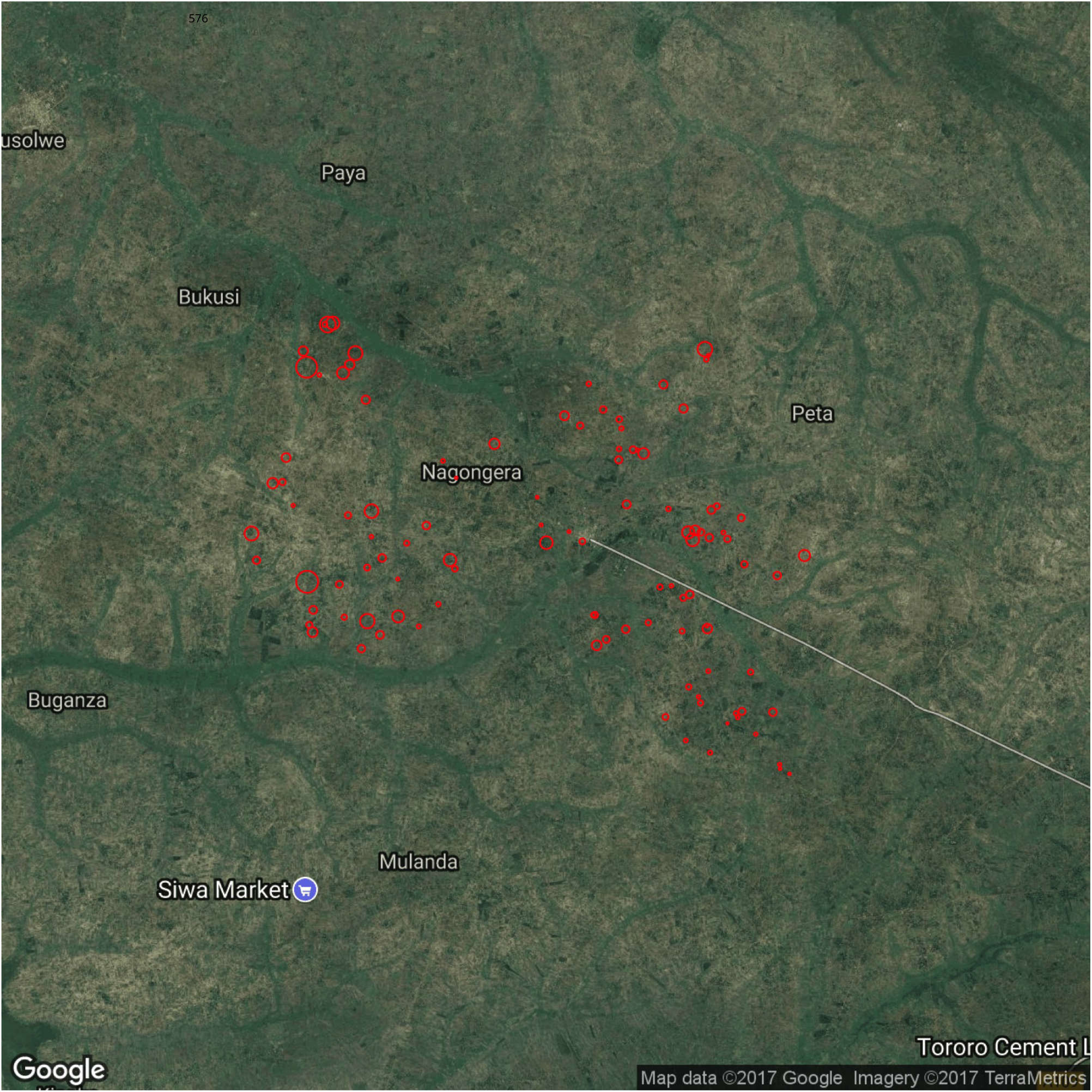
Sampled households in Nagongera subcounty, Tororo District between October 2011 and September 2016, located in south-eastern Uganda on the Kenyan border. Each red circle denotes a household and the size ofthe circles denotes the overall biting propensities over the entire duration of surveillance. This is the site with highest malaria transmission in our study.

## Supplementary File 2: Detailed description of simulation analyses

### Model description

Generalized linear models (McCullagh and Nelder, 1989) are employed in this study to model mosquito counts. The response variable (mosquito counts) is assumed to be generated from a probability distribution from the exponential family, and the mean ofthe distribution is related to the independent variables through a link function. Here we adopt a Bayesian framework to estimate model parameters, using the integrated nested Laplace approximation (INLA) approach (Rue et al., 2009).

The mosquito counts at the study sites in Uganda consist of a large proportion of zero values; 70% in Jinja, 54% in Kanungu, and 21% in Tororo. We therefore consider specific probability models that are capable of handling excess zeros while modeling non-zero counts properly (Cohen, 1963; Hilbe, 2014), namely zero-inflated (Lambert, 1992; Cameron and Trivedi, 2013) and hurdle (Cragg, 1971; Mullahy, 1986) models. These two models can be viewed as mixture models with subtle differences between them. By fitting zero-inflated Poisson (ZIP), zero-inflated negative binomial (ZINB), Poisson hurdle (PH), and negative binomial hurdle (NBH) regression models to the mosquito counts, we estimate household biting propensities **(*ω*)**, seasonal signal **(*S*)**, and environmental noise and measurement error **(*e*)** at each study site.

A strong seasonal pattern is observed in the mosquito counts for the three sites. It is thus necessary to perform seasonal adjustment (Sims, 1974) by removing the seasonal component of the time series data in order to understand the components ofthe underlying trends in mosquito biting. The seasonal signal in the mosquito counts across years is captured using two main categories of temporal smoothing techniques in order to discover the optimal way for capturing seasonal trend: (1) a Gaussian smoothing kernel (Wand and Jones, 1995) that adjusts for seasonality prior to model fitting; and (2) Bayesian prior distributions for temporally structured random effects.

With respect to technique (1), seasonal smoothing is implemented using the ‘KernSmooth’ package in R (Wand and Ripley, 2006), which selects a bandwidth for kernel density estimation using a direct plug-in approach. The estimated function using a Gaussian kernel is smooth, and the level of smoothness is controlled by a single parameter, bandwidth *h*; expressed below,

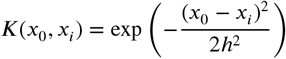

With respect to technique (2), we explore several prior distributions which accommodate temporally varying smoothing functions for modeling time effects ofthe observed counts. The available time points (sampling days) are modeled as structured random effects, ensuring that contiguous periods are likely to be similar, but allowing for flexible shapes in the evolution curve. First– and second-order random walks (RW1, RW2), and autoregressive processes of order 1 and order 2 (AR1, AR2) are considered here (Sørbye and Rue, 2014) and are implemented using R-INLA (Lindgren and Rue, 2015). To summarize, the following seasonal smoothing techniques are undertaken for each ofthe models,

S1: A Gaussian kernel is used to smooth the raw data and the model is fitted to the smooth data;

S2, S3, S4, and S5: A RW1, RW2, AR1, and AR2 prior is imposed on sampling days (Day) in the model for the non-smooth data, respectively.

#### Zero-inflated models

A zero-inflated model (Lambert, 1992; Cameron and Trivedi, 2013) is a mixture of a point mass at zero and a count distribution. In this model, zero counts may arise from one ofthe two data generating processes, either by chance (i.e., a process that generates nonzero counts – condition is present but a zero is recorded by “mistake” in some cases as they may not be detected), or by definition (i.e., a process that generates only zeros – a zero is recorded since condition is absent) (Arab, 2015). The former is also known as “sampling” zeros whereas the latter is known as “structural” or true zeros.

Here we consider two count distributions for the zero-inflated model. For a zero-inflated Poisson (ZIP) distribution, we can write

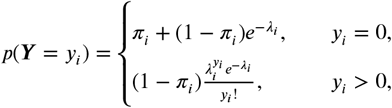

for the mosquito counts ***Y*** = {*y*_1_, *y*_2_,…, *y_n_*}, with probability 0 ≤ *π_i_* ≤ 1, and *λ_i_* ≥ 0 is the expected count ofthe Poisson distribution. The mean ofthe ZIP distribution is *E*(*Y_i_*) = (1 – *π_i_*)*λ_i_* and the variance is var(*Y_i_*) = *λ_i_*(1 – *π_i_*)(1 + *λ_i_π_i_*). The zero-inflated negative binomial (ZINB) distribution can be written as

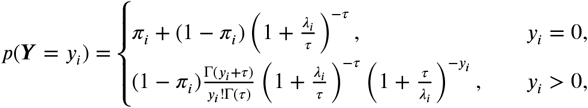

where *τ* > 0 is a shape parameter which quantifies the amount of overdispersion. The mean and variance ofthe ZINB distribution are ***E*(*Y_i_*)** = (1 – *π_i_*)*λ*_i_ and var**(Y**_i_) = *λ*(1 – *π_i_*)(1 + *π_i_λ_i_* + *λ_i_*/*τ*), respectively.

#### Hurdle models

A hurdle model (Cragg, 1971; Mullahy, 1986) consists of two components – a point mass at zero and a distribution that generates non-zero counts. The first component is a binary component that generates zeros and ones (here “ones” correspond to non-zero values in data) and the second component generates non-zero values from a zero-truncated distribution. Note that hurdle models have a general interpretation and the “hurdle” may be any value other than zero. The most widely used hurdle models are those with the hurdle value at zero (Hilbe, 2014). All zeros in the hurdle model are assumed to be “structural” zeros, i.e., they are generated from a single process, and are observed since the condition is absent.

We explore two zero-truncated count distributions for the hurdle model specification. A Poisson hurdle (PH) model for the observed counts ***Y*** = {*y*_1_, *y*_2_,…, *y_n_*} can be described as the mixture of a point mass at zero with probability *π_i_* and a zero-truncated Poisson distribution with probability l – *π_i_*:

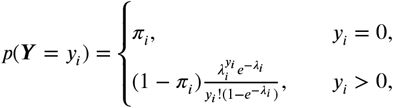

where *λ_i_* is the mean ofthe untruncated Poisson distribution. A negative binomial hurdle (NBH) distribution (Pohlmeier and Ulrich, 1995; Arulampalam and Booth, 1997; Saffari et al., 2012) is given by

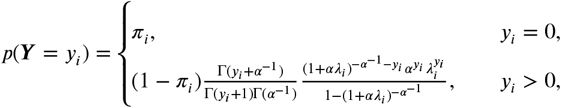

where *α*(≥ 0) is a dispersion parameter that is assumed not to depend on covariates.

### Parameter estimation

Note that *p***(*Y*** = *y_i_*) is a function of *π_i_* and *λ_i_* in the ZIP, ZINB, PH, and NBH regression models. The parameters *π_i_* and *λ_i_* can be modeled as a function of a set of explanatory variables. A logistic regression with a logit link function is used to model *π_i_*, as it describes a binomial process. A log link function is used to model the dependence of *λ_i_* on a different (or same) set of covariates. The log link function ensures that the estimated *λ_i_* will not be negative, regardless of parameter values. In our case, we model the dependence of *π_i_* and *λ_i_* on the same set of explanatory variables.

For each ofthe three sites, the ZIP, ZINB, PH, and NBH regression models relate *λ_ij_*, the expected count for household *j* on day *i*, to covariates on a logarithmic scale. Using the same set of covariates, the probability *π_ij_* is modeled using a logit link function. The covariates of interest is ID, the household identifiers. The model corresponding to S1 is

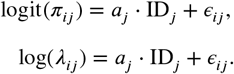

The model corresponding to S2, S3, S4, and S5 is

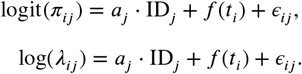

Here *a_j_* quantifies the effects of household biting. The random effects, *∈_ij_*, are assumed to be independent and identically distributed, and *t_i_* are the temporally structured random effects assigned one ofthe four Bayesian seasonal smoothing priors. From these estimates, we obtain the household biting propensities, ***ω*** = exp(*a_j_*) and the seasonal signal, ***S*** = exp(*f*(*t_i_*)). The environmental noise and measurement error, ***e*** = {*e_ij_*}, accounts for additional variation among mosquito counts that is not accounted for by the covariates or by Poisson (random) variation about the means ***ωS***. The distribution ofthe noise term, ***e***, was obtained by modeling *∈_ij_* with a Poisson-Gamma mixture process. Other mixture process such as a Poisson-lognormal distribution (Clayton and Kaldor, 1987) may also be appropriate. Put simply, ***e*** accounts for the remaining variability in mosquito counts which is not captured by household biting propensities, seasonality, and other relevant covariates.

### Simulation study

The simulation study was designed to determine which method was the most appropriate for disentangling the signal and noise in mosquito counts. The data generated consisted of sampling days, mosquito counts, and household identifiers. The simulated data were observed to be noisy across sampling days, with a seasonal trend, and were overdispersed (the variance larger than the mean). We considered six pseudo-datasets which differed in the true distribution of household biting propensities, namely D1, D2, D3, D4, D5, and D6. The pseudo-data were generated based on the following procedures. For household *j* and day *i*, the mosquito counts followed a Poisson distribution: *y_ij_* ~ **Poisson(*ω_j_S_i_e_ij_*)**. True biting propensities were simulated for each household drawn from either a Gamma distribution or a log-normal distribution: *ω_j_* ~ ***Γ*(*α,β*)** or *ω_j_* ~ ln ***N*(*μ, σ***^2^). The seasonal signal was defined to follow the multiplication of several sigmoid functions, ***S**_i_* = 5(*pqrst*)^1.5^, where

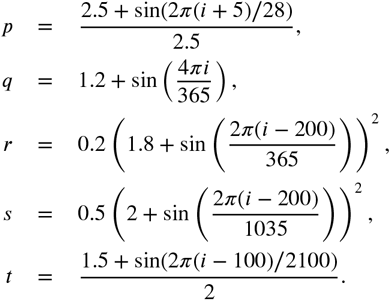

The environmental noise and measurement error followed a Gamma distribution: *e_ij_* ~ Γ(*α* = 1.3, *β* = 1.3). For a variety of methods discussed above, we estimated the household biting propensities (***ω**_P_*), seasonal signal (***S**_p_*), and environmental noise and measurement error (*e_P_*) for all six pseudo-datasets, and compared them to the truth.

The performance of various methods in disentangling the effects of ***ω**_P_, **S**_P_*, and ***e**_P_* for the pseudo-datasets was of primary interest here (note that the subscript ***P*** denotes pseudo-datasets). For simplicity, our description ofthe results focuses on the results of pseudo-dataset D3, but we stress that the results of other pseudo-datasets are similar. The ZINB model was the best model in recovering true parameters for the distribution of the household biting propensities, ***ω**_P_*, for all pseudo-datasets (Figures A1 (a) and A2) as its estimated parameters were the closest to the true parameter values in all settings when compared among all models. The use of S1 for seasonal smoothing in all models led to poor estimation of parameters, whereas S2, S3, S4, and S5 performed much better than S1 and showed comparable performance in recovering the true parameters of ***ω**_P_*

With respect to recovering the seasonal signal, the ZINB model (with either S2, S3, S4, or S5) produced the smallest root mean square error (RMSE) computed between the true seasonal signal, ***S**_P_*, and the reconstructed 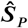. Different smoothing techniques yielded RMSEs with little differences between them for the ZIP and ZINB models, but with larger differences observed between themfor the PH and NBH models. Note that the RMSE produced by S1 was not affected by the choice of model because the Gaussian kernel smoothing was undertaken prior to model fitting. Again, Figures A1 (b) confirmed that the seasonal signal reconstructed using the ZINB S3 model was the closest to the truth.

For the identification ofthe seasonal signal, the use of a Gaussian smoothing kernel on raw mosquito data was not recommended because it often seemed to produce over-smoothed seasonal estimates. Seasonal smoothing should be undertaken within the model using a seasonal smoothing prior distribution for the temporal effects instead. We discovered that the various seasonal smoothing priors considered in the simulation study had performed equally well in recovering the true seasonal signal and that the final choice of the smoothing prior could be determined using a model selection criterion, such as the Watanabe-Akaike information criterion (WAIC) (Watanabe, 2010).

With respect to disentangling the noise from other signals, again, the ZINB model outperformed all other models (Figure A1(c)), by producing the smallest RMSE between the true ***e**_P_* and the estimated 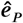. In short, the ZINB model was shown to be the most robust method in disentangling signals and noise in mosquito abundance across all settings in this simulation study. The true distribution ofthe mosquito counts is compared against the distribution simulated from the ZINB model for each ofthe pseudo-datasets (Figure A3). Clearly, the simulated distributions of mosquito counts closely resembled the true distributions of counts for all datasets, which suggested that our method worked efficiently for various types of distributions of data.

**Figure A1.**
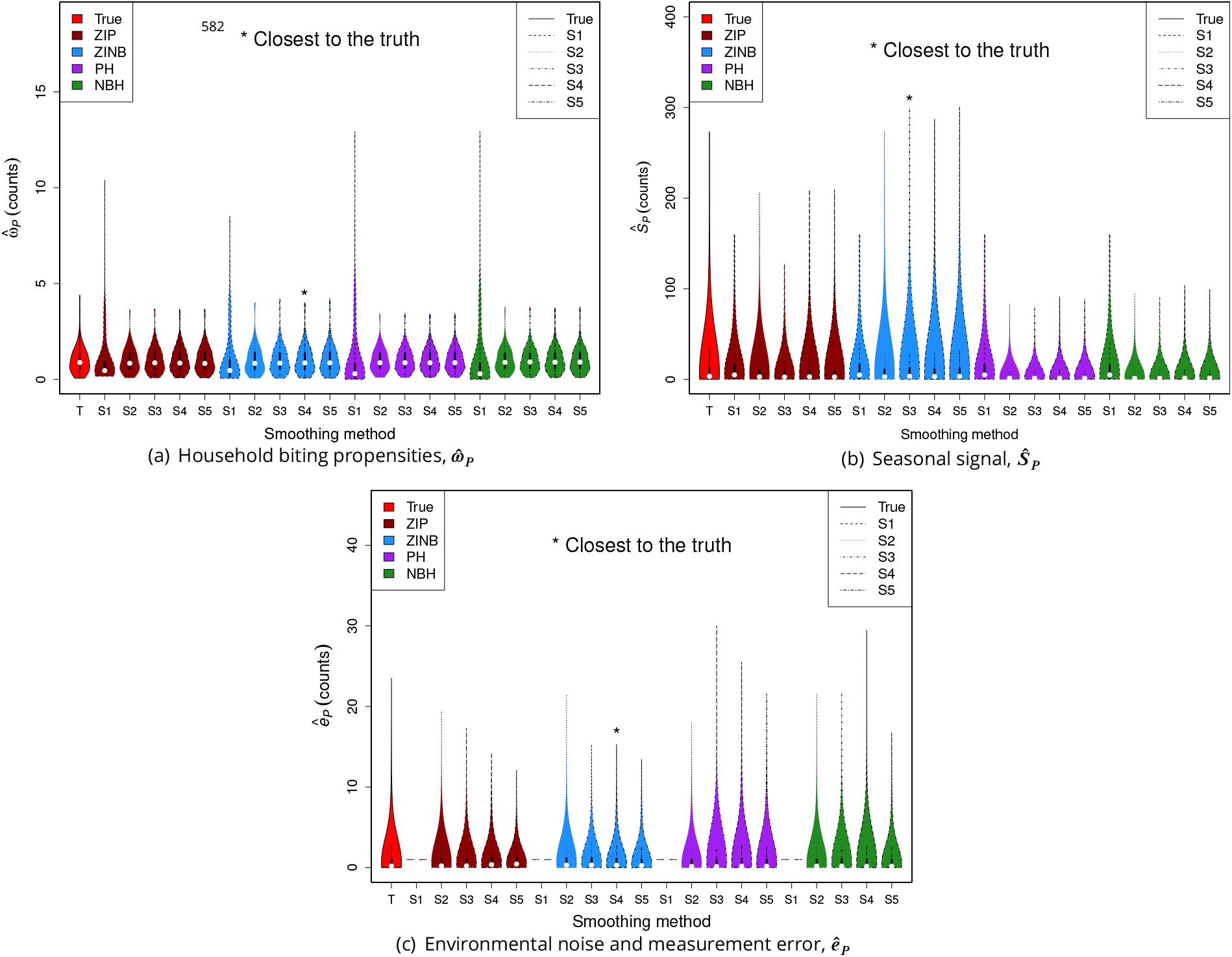
Simulation study: The estimated 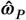, 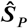, and 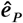 for pseudo-dataset D3 using various models and smoothing techniques. The method that produces estimates closest to the truth is marked with *. Note that 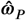 and 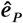 were normalized to have a mean of 1.

**Figure A2.**
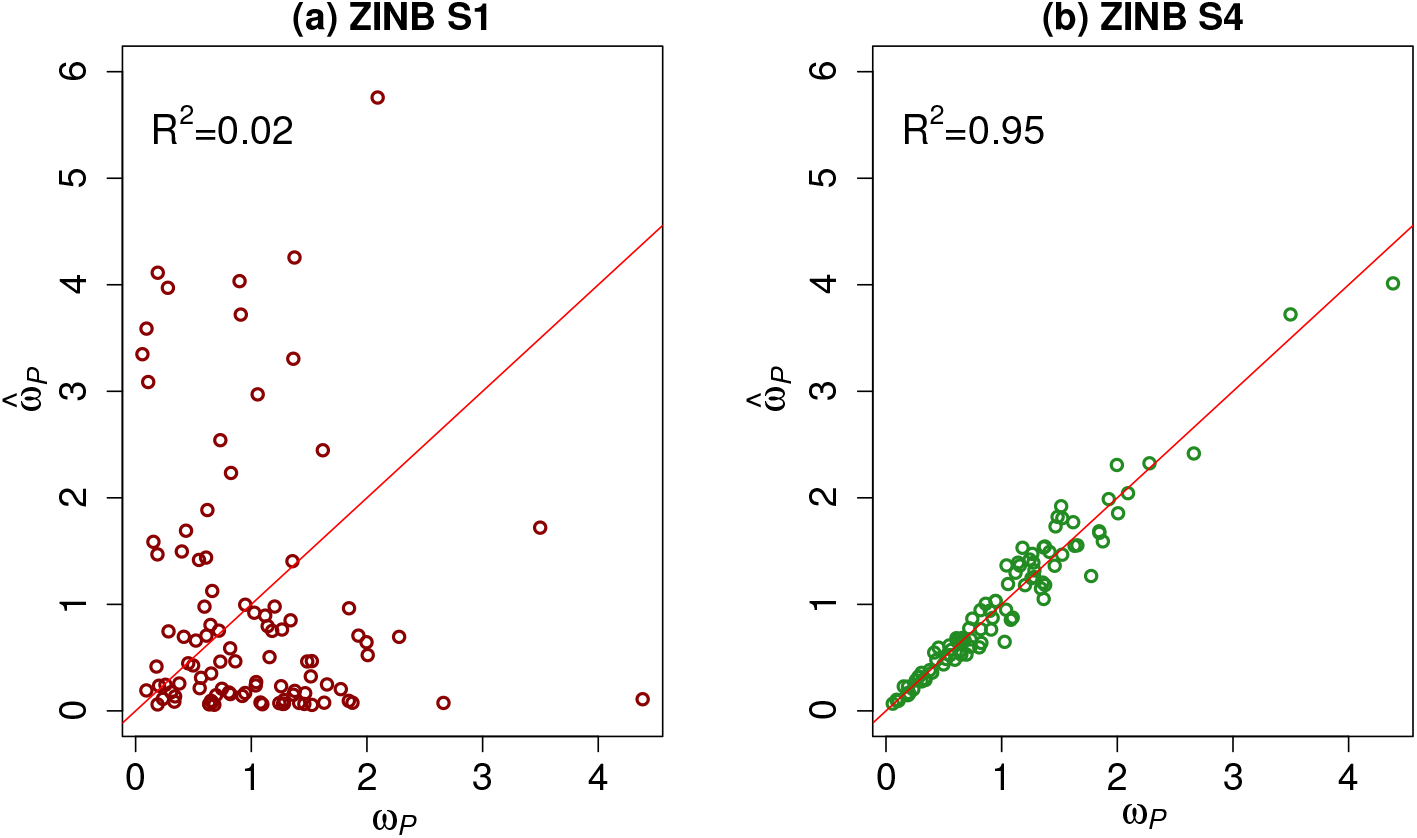
Simulation study: The estimated household biting propensities, *ω*_P_, against the true *ω_P_* for pseudo-dataset D3 using the best model, i.e. the ZINB model. (a) S1 produced the worst fit when *ω_P_* is fitted against *ω_P_* on a simple linear regression; coefficient of determination, *R*^2^, is close to 0. (b) S4 produced the best fit when *ω_P_* is fitted against *ω_P_* on a simple linear regression; *R*^2^ is close to 1.

**Figure A3.**
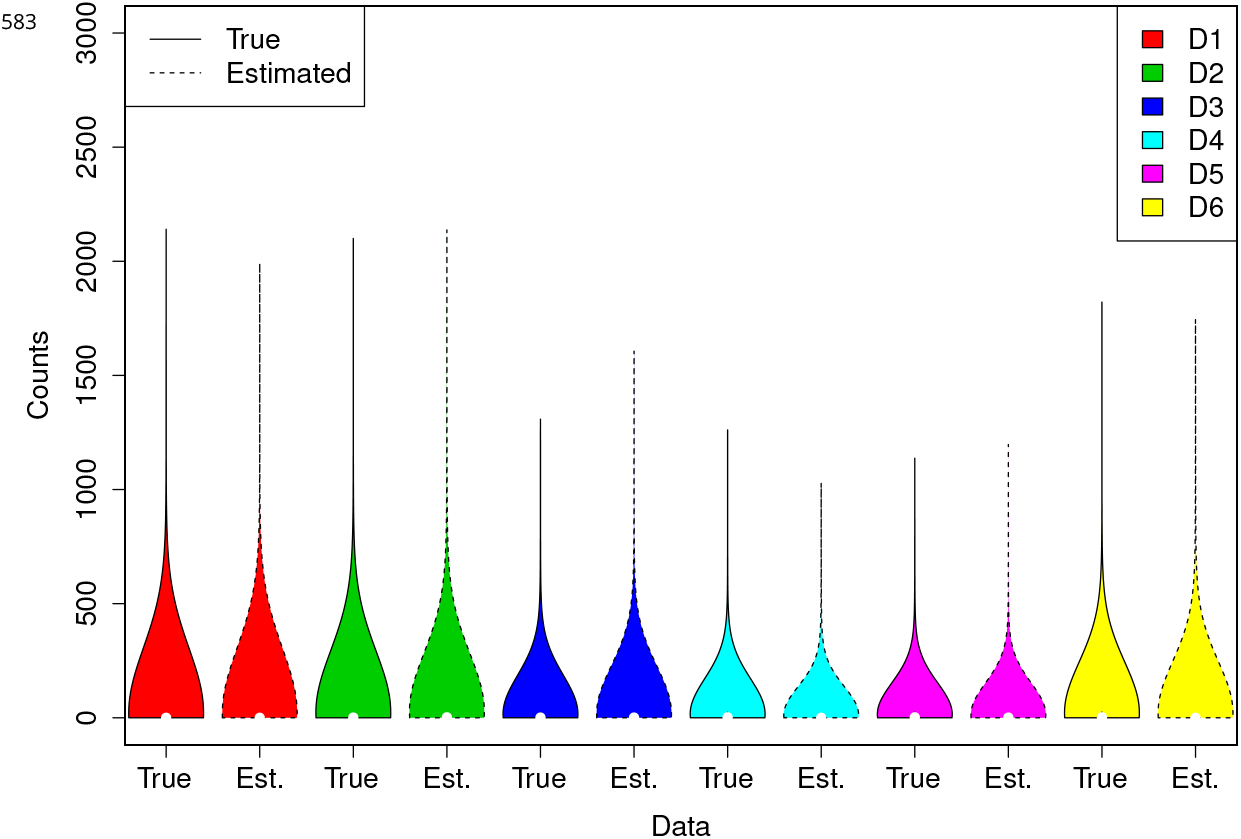
Simulation study: The true distribution and the estimated distribution of mosquito counts for all six pseudo-datasets. The estimated distributions were simulated using the best estimates of *ω*_P_, 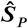, and *ê_P_* from the ZINB model for each dataset, respectively. The simulated distributions of counts closely resembled the true distributions of counts for all datasets.

